# Analyzing genomic alterations involved in fluoroquinolone-resistant development in *Staphylococcus aureus*

**DOI:** 10.1101/2023.02.26.530158

**Authors:** Thuc Quyen Huynh, Van Nhi Tran, Van Chi Thai, Hoang An Nguyen, Ngoc Thuy Giang Nguyen, Navenaah Udaya Surian, Swaine Chen, Thi Thu Hoai Nguyen

**Affiliations:** School of Biotechnology, International University, Vietnam National University of Ho Chi Minh City, Vietnam; Research Center for Infectious Diseases, International University, Vietnam National University of Ho Chi Minh City, Vietnam; Genome Institute of Singapore, Singapore

**Keywords:** Fluoroquinolone, multidrug resistance, *Staphylococcus aureus*, whole genome sequencing

## Abstract

**Aim:** Recently, the rise in Staphylococcal infection incidence accompanied by a rise of antibiotic-resistant strains is a major threat to public health. In this study, mechanisms leading to the occurrence of high-level multidrug-resistant (MDR) *Staphylococcus aureus (S. aureus)* strains after fluoroquinolone (FQ) exposure were investigated.

**Methodology:** Serially exposing *S. aureus* ATCC 29213 to ciprofloxacin (CIP), ofloxacin (OFL), or levofloxacin (LEV) at sub-minimum inhibitory concentrations (sub-MICs) for 12 days was performed to obtain *S. aureus* -1 strains and culturing for another 10 days without antibiotics to obtain *S. aureus*-2 strains. The genomic alterations in FQ-exposed strains were reached using whole genome sequencing and target sequencing. The expressions of efflux-related genes, alternative sigma factors, and genes involved in FQ resistance were evaluated using RT-qPCR.

**Results:** After serial FQ exposure, we observed a strong and irreversible increase of MICs to all applied FQs, i.e 32 to 128 times in all *S. aureus*-1 and remained 16 to 32 times in all *S. aureus*-2. WGS indicated 10 significant mutations including 2 deletions, 1 insertion, and 7 missense mutations that occur in all *S. aureus*-1 and -2 but not in initial strain. The FQ target, GrlA, was also mutated (R570H) in all *S. aureus*-1 and -2 which can partly explain the development of FQ resistance over the FQ exposure. Besides, FQ exposure also resulted in overexpression of genes encoding for (1) efflux pumps and their regulator (*norA, norB, norC*, and *mgrA*); (2) alternative sigma factors (*sigB* and *sigS*); (3) acetyltransferase (*rimI*); (4) methicillin resistance (*fmtB*); and (5) hypothetical protein BJI72_0645.

**Conclusion:** The mutations occurred in the FQ-target sequence were associated with high-level FQ resistance while the activation of efflux pump systems and post-translational proteins played an important role in the emergence of MDR in *S. aureus*.

**Author summary:** Antimicrobial resistance is a major public health problem worldwide. Multiple studies have been performed to understand how bacteria develops resistance during the antibiotic therapy *in vitro* and *in vivo*. Here we revealed how *Staphylococcus aureus*, a stubborn human pathogen, changed its genome and expression of important genes in responding with sub-MIC exposure to flouroquinolone antibiotics. Mutations were found in the target of flouroquinolones such as GrlA (R570H) and interestingly in some hypothetical regions which may be important for gene expression regulation. We have observed an marked overexpression of genes encoding for (1) efflux pumps and their regulator (*norA, norB, norC, and mgrA*); (2) alternative sigma factors (*sigB and sigS*); (3) acetyltransferase (*rimI*); (4) methicillin resistance (*fmtB*); and (5) hypothetical protein BJI72_0645 in all exposed strains. Importantly, the expression change still remained when the bacteria were no longer exposed to the antibiotics. This study is important to understand response of *S. aureus* to flouroquinolone and how it obtains the resistance phenotype under antibiotic exposure.

## Introduction

*Staphylococcus aureus (S. aureus)* is a common extracellular pathogen with the ability to cause a wide range of infections ranging from mild (skin infections, and soft tissues) to serious (life-threatening infections, such as pneumonia, osteomyelitis, endocarditis, toxic shock syndromes…) [1]. The rates of multidrug resistance in *S. aureus* have been increasing considerably in recent years, leading to serious consequences, including treatment failure, treatment cost consumption, and protracted therapy [2,3].

Fluoroquinolone (FQ) is a unique class of synthetic broad-spectrum antibiotics that has been applied for the treatment of a wide variety of community and nosocomial infections [4]. Unfortunately, the broad use of these drugs for clinical treatment has led to a steadily increasing number of FQ-resistant bacterial strains, with rates of resistance that vary with both organism and geographic region [5]. The proportion of Gram-positive cocci resistance to these drugs also appears to be increasing worldwide, especially for *S. aureus* [6].

The resistance of *S. aureus* to FQs results from mutations in quinolone resistance-determining regions (QRDRs) of topoisomerase IV (*grlA/B*) and/or DNA gyrase (*gyrA/B*) [7,8] and the over-expression of efflux pumps systems transporting the antibiotics out of the cells [9,10]. It is assumed that some of the specific mutations in the topoisomerase IV and DNA gyrase gene identified from clinical strains were involved in FQ resistance [8,11,12]. Although these alterations appear to decrease their affinity to FQs [7,13–15], it is difficult to analyze the contribution of each mutation to the resistance phenotype.

Thus, the aim of this study was to investigate the mechanism of antibiotic resistance development in *in vitro* selected FQ-resistant *S. aureus* ATCC 29213. To achieve this, we performed whole genome sequencing of *in vitro* selected FQ-resistant *S. aureus* strains as well as target sequencing of important genes. Additionally, the expression of efflux-related genes, alternative sigma factors, and genes involved in FQ resistance were evaluated using RT-qPCR were also evaluated.

## Methods

### Selection of FQ-resistant strains

*S. aureus* ATCC 29213 (initial *S. aureus*) was cultured in Mueller-Hinton Broth (MHB) that contained the continuous presence of fluoroquinolone (FQ) either ciprofloxacin (CIP), ofloxacin (OFL) or levofloxacin (LEV) with concentrations corresponding to half of the minimum inhibitory concentration (MIC) value [16]. The experiment was repeated until no increase in MIC to the FQ used in exposure was observed. At the endpoint, the obtained CIP-, LEV-, and OFL-exposed *S. aureus* (*S. aureus*-1) strains were sent to NK Biotechnology Company (HCM, Vietnam) for *16S rRNA* sequencing. These selected resistant strains were sub-cultured for another 10 days in antibiotic-free medium with daily examination of MIC in order to obtain CIP-, LEV-, and OFL-revertant *S. aureus* (*S. aureus*-2) strains. Repetitive sequence-based PCR (Rep-PCR) amplification was established to confirm the genetic relatedness of initial *S. aureus* ATCC 29213 and those of selected strains. The test was adapted from Vito *et al*. [17]. The strains at day 12^th^ of FQ exposure (*S. aureus*-1 including CIP-, OFL- and LEV-1) and the strains at day 10^th^ of antibiotic-free culture (*S. aureus*-2 including CIP-, OFL- and LEV-2) were then used for other experiments.

### Antimicrobial susceptibility testing

The micro-dilution method on 96-well plates, instructed by EUCAST guidelines (Eucast.org, version 11.0), was applied on seven bacterial strains (initial *S. aureus, S. aureus*-1 and -2) to determine the MIC value of the strains to CIP, LEV, OFL, moxifloxacin, nalidixic acid, ampicillin, amoxicillin, chloramphenicol, cefalexin, doxycycline, erythromycin, lincomycin, oxacillin, and tetracycline. At the same time, MICs of the strains were also measured under the presence of an efflux pump inhibitor, reserpine (Sigma-Aldrich, USA), at a concentration of 20 mg/L.

### Whole genome sequencing

DNA extraction was carried out using the GeneJET Genomic DNA Purification Kit (Thermo Scientific, USA) according to the manufacturer’s instructions. Sequencing libraries were prepared using the TruSeq Nano DNA LT Library Prep Kit (Illumina, Singapore). Pooled libraries were then sequenced using an Illumina NextSeq 500 sequencer with 2 × 151 bp reads. The resulting FASTQ files were mapped to the ATCC 29213 genome (Genbank GCF_001879295.1) using bwa (version 0.7.10) (PMID 19451168); indel realignment and SNP (single nucleotide polymorphism) calling were performed using Lofreq* (version 2.1.2) with default parameters (PMID 23066108).

### Quinolone-resistance determining regions (QRDRs) target sequencing

The genomic DNA of *S. aureus* strains (Initial *S. aureus* ATCC 29213, *S. aureus* strains at day 4^th^, 6^th^, 8^th^, 10^th^ of CIP-, LEV- and OFL-exposure, CIP-, LEV-, OFL-resistant strain-1 and -2) were extracted and then, utilized for QRDRs (DNA gyrase -*gyrA* and *gyrB*, and topoisomerase IV -*grlA* and *grlB*) amplification. Each of 50 µl PCR reaction was prepared with instructions from the *TopTaq Master Mix* Kit (Qiagen, Germany). The amplification profiles and primers were described previously [8,18]. The PCR products were finally electrophoresed and subjected to sequencing (Macrogen, Korea).

### Expression evaluation of efflux-related genes, alternative sigma factors, and genes involved in FQ resistance

Total RNA extraction and cDNA Synthesis of seven *S. aureus* strains were performed using Monarch® Total RNA Miniprep Kit (NEB, UK) and SensiFAST cDNA Synthesis Kit (Bioline, UK), according to the manufacturer’s instructions. Each real-time qRT-PCR reaction consisted of 20 μL of reagents, including Luna® Universal qPCR Master Mix (NEB, UK), one primer pair (***cDNA template***, and nuclease-free water. The thermal cycle of the RT-qPCR machine was set up based on NEB #M3003 instruction manual (NEB, UK). Transcription values (Ct) are analyzed as described in [19]. All experiments are done in triplicate.

**Table 1**; Reference gene: *16S rRNA*), cDNA template, and nuclease-free water. The thermal cycle of the RT-qPCR machine was set up based on NEB #M3003 instruction manual (NEB, UK). Transcription values (Ct) are analyzed as described in [19]. All experiments are done in triplicate.

**Table 1.**
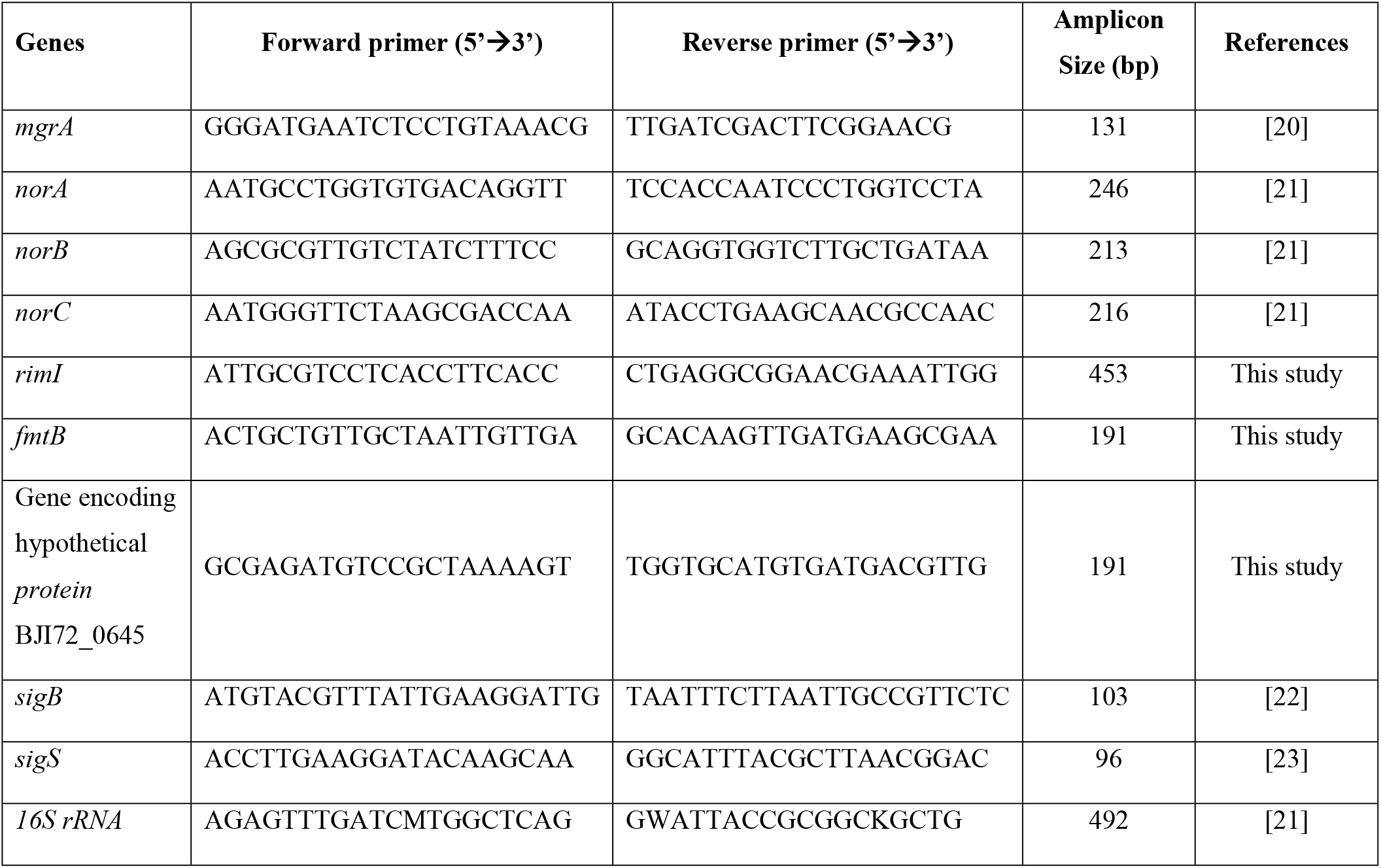
Primers for qRT-PCR analysis.

## Results

### Serial exposure to FQs leading to FQ-resistant phenotype

After being serially exposed to FQs, *S. aureus* ATCC 29213 turned from FQ-sensitive to FQ-resistant phenotype. The MIC values of the FQ-exposed strains reached peaks at day 8^th^-10^th^ of exposure (***Fig 1***), which was 128 times higher than the MIC of initial strain for CIP, and 32 times higher for LEV and OFL. No further increase in MIC values was observed after the peaks even if these strains were continuously exposed to the antibiotics. Impressively, these FQ-exposed strains did not revert to antibiotic-sensitive phenotype when being cultured in an antibiotic-free medium for 10 days (remained 8-32 times higher than initial strain). Gram staining, *16S rRNA*, and Rep-PCR sequencing results proved no contamination during the antibiotic exposure process (data not shown). In addition, molecular genotyping results showed that *S. aureus*-1 and -2 strains were identical to initial *S. aureus* ATCC 29213.

**Fig 1.**
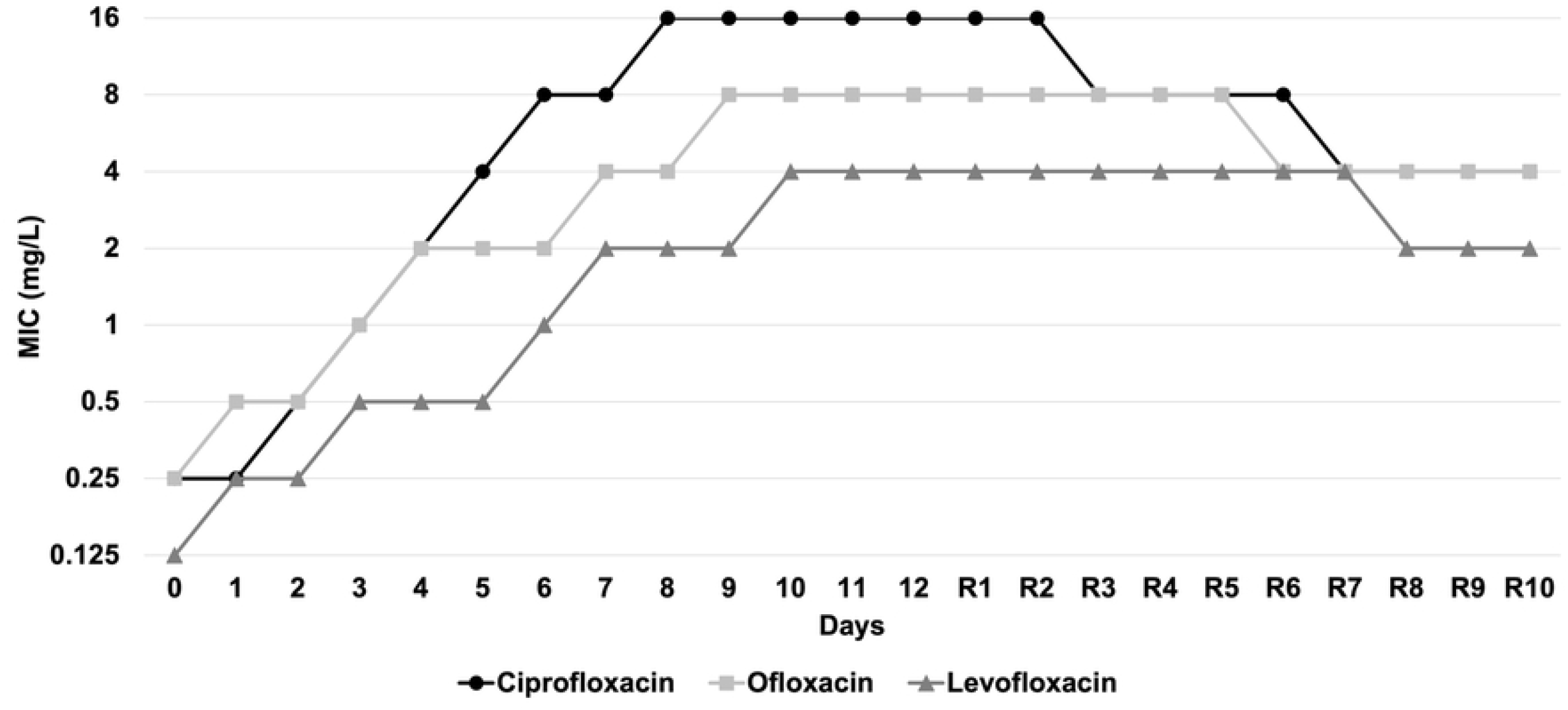
MIC values of *S. aureus* during sub-MIC exposure to fluoroquinolones (FQs). The initial *S. aureus* was exposed to FQs (ciprofloxacin, levofloxacin, and ofloxacin) for 12 days for obtaining FQ-resistant *S. aureus* strains. Day 0: MIC; day 1-12: 12 FQ exposed days. These 12 days of exposed *S. aureus* (CIP, LEV, and OFL-resistant *S. aureus*) were continuously sub-cultured for another 10 days (R1-R10) in an antibiotic-free medium with the daily examination of MICs.

### Susceptibility of *S. aureus* strains to different antibiotics

The FQ-exposed *S. aureus* ATCC 29213 turned into a multidrug-resistant phenotype, which was not only resistant to other FQs but also to other antibiotics of unrelated groups (***Table 2***). The increase in MICs of *S. aureus-*1 and -2 strains were as followings: ampicillin (8-16, 4-16 times), amoxicillin (64-128 times), cefalexin (4-128 times), doxycycline (16-32 times), erythromycin (64-128 times), lincomycin (64-256, 64-128 times) and oxacillin (16-64, 16-32 times). Interestingly, the exposed strains remained sensitive to moxifloxacin (0.25-0.5 mg/L) and unchanged to chloramphenicol and tetracycline. Subsequent cultures of *S. aureus-*1 strains in an antibiotic-free medium only resulted in minor effects on their susceptibility. Most MIC values of *S. aureus-*2 strains decreased only 2 to 4 times compared to *S. aureus-*1 strains. Under the presence of reserpine, *S. aureus* ATCC 29213 was not affected while susceptibility of *S. aureus*-1 strains was reduced by 2-8 folds for some antibiotics such as ampicillin, amoxicillin, doxycycline, erythromycin, and lincomycin (***S1 Table***).

**Table 2.** Antibiotic susceptibility of *S. aureus* strains.

| Antibiotics | *S. aureus* strains |  |  |  |  |  |  |
| --- | --- | --- | --- | --- | --- | --- | --- |
|  | Initial | CIP-1 | CIP-2 | LEV-1 | LEV-2 | OFL-1 | OFL-2 |
| ***Fluoroquinolone*** | MIC (mg/L) |  |  |  |  |  |  |
| Ciprofloxacin | 0.125 | 16 | 4 | 8 | 2 | 4 | 2 |
| Levofloxacin | 0.125 | 4 | 1 | 4 | 2 | 8 | 4 |
| Ofloxacin | 0.25 | 4 | 2 | 16 | 4 | 8 | 4 |
| Moxifloxacin | 0.0625 | 0.25 | 0.25 | 0.5 | 0.5 | 0.5 | 0.25 |
| Nalidixic acid | 16 | 128 | 64 | 128 | 64 | 128 | 64 |
| ***Unrelated antibiotics*** |  |  |  |  |  |  |  |
| Ampicillin | 16 | 256 | 256 | 128 | 64 | 128 | 64 |
| Amoxicillin | 0.5 | 32 | 32 | 64 | 32 | 64 | 64 |
| Cefalexin | 0.5 | 4 | 4 | 64 | 64 | 8 | 8 |
| Chloramphenicol | 16 | 32 | 32 | 16 | 16 | 32 | 16 |
| Doxycycline | 0.125 | 4 | 4 | 4 | 4 | 2 | 2 |
| Erythromycin | 0.25 | 32 | 16 | 16 | 16 | 32 | 32 |
| Lincomycin | 0.5 | 32 | 32 | 64 | 32 | 128 | 64 |
| Oxacillin | 0.5 | 16 | 16 | 8 | 8 | 32 | 16 |
| Tetracycline | 8 | 16 | 16 | 16 | 16 | 8 | 8 |

### Whole genome sequencing of *S. aureus* ATCC 29213, *S. aureus*-1 and *S. aureus*-2 strains

Whole genome sequencing was used to identify mutations associated with the changes in resistance (***S2 Table***). Overall, a total of 42 positions where at least one strain carried a variant relative to initial strain were discovered. Of these 42 variant positions, 10 were common in all strains sequenced, indicating differences in our parental strain from the publicly available genome sequence. Among 32 remaining variant positions, 24 were SNPs, 6 were deletions, and 2 were insertions. A total of 11 variants were seen in all 6 of the *S. aureus*-1 and *S. aureus*-2 strains relative to our re-sequenced ATCC 29213 (***Table 3***). Of these 11 SNPs, 4 were found in predicted protein-coding genes: an amino acid deletion in SACOL0573 (encoding a PIN domain-containing protein); an amino acid insertion in *rimI* (an alanine acetyltransferase); a missense mutation in *grlA*, encoding DNA topoisomerase IV subunit A, and a synonymous mutation in *hslO*, which encodes heat shock protein Hsp33. The remaining 7 variants located in the upstream region of *norA* and gene coding hypothetical protein BJI72_0645 were not found in annotated protein-coding sequences.

**Table 3.** The list of single nucleotide polymorphisms (SNPs), variants, and amino acid changes found in *S. aureus*-1 (CIP-1, OFL-1, and LEV-1) and *S. aureus*-2 (CIP-2, OFL-2, and LEV-2) strains but not in *S. aureus* ATCC 29213.

| SNP/variant and<br>Genebank Accession | Amino acid changes | Systematic and<br>trivial gene<br>name | Protein encoded | Mutations in samples |  |  |  |  |  |  |
| --- | --- | --- | --- | --- | --- | --- | --- | --- | --- | --- |
|  |  |  |  | *S. aureus*<br>ATCC<br>29213 | CIP-1 | OFL-1 | LEV-1 | CIP-2 | OFL-2 | LEV-2 |
| G5009GAAC<br>MOPB01000035.1 | S33_S34insC | BJI72_1850<br>rimI | Alanine<br>acetyltransferase | . | √ | √ | √ | √ | √ | √ |
| TATCTAGATGA<br>TG4318T<br>MOPB01000016.1 | G309_T315delinsC | BJI72_0480<br>SACOL0573 | PIN domain<br>containing protein | . | √ | √ | √ | √ | √ | √ |
| A13766G<br>MOPB01000012.1 | No annotated<br>protein | - | - | . | √ | √ | √ | √ | √ | √ |
| A13775T<br>MOPB01000012.1 | No annotated<br>protein | - | - | . | √ | √ | √ | √ | √ | √ |
| A13777C<br>MOPB01000012.1 | No annotated<br>protein | - | - | . | √ | √ | √ | √ | √ | √ |
| A13778T<br>MOPB01000012.1 | No annotated<br>protein | - | - | . | √ | √ | √ | √ | √ | √ |
| T13781A<br>MOPB01000012.1 | No annotated<br>protein | - | - | . | √ | √ | √ | √ | √ | √ |
| **T13785G**<br>**MOPB01000012.1** | **No annotated protein** | - | - | . | √ | √ | √ | √ | √ | √ |
| **CG13787C**<br>**MOPB01000012.1** | **No annotated protein** | - | - | . | √ | √ | √ | √ | √ | √ |
| **C32411A**<br>**MOPB01000008.1** | **Synonymous mutation** | **BJI72_0464**<br>**hslO** | **heat shock protein**<br>**Hsp33** | . | √ | √ | √ | √ | √ | √ |
| CATAGGCTTGTT<br>22457C<br>MOPB01000021.1 | N3181delinsR | BJI72_1235<br>Ebh | Hyperosmolarity<br>resistance protein Ebh | . | . | √ | √ | . | √ | √ |
| T86957C<br>MOPB01000021.1 | Synonymous<br>mutation | BJI72_1284<br>lpl | lipoprotein | . | . | √ | √ | . | √ | √ |
| CTT26104C<br>MOPB01000037.1 | K1283delinsRG | BJI72_1954<br>fmtB | methicillin resistance<br>protein FmtB | . | . | √ | √ | . | √ | √ |
| C93466T<br>MOPB01000004.1 | A318T | BJI72_0139<br>ThlA | acetyl-CoA<br>acetyltransferase | . | √ | . | . | √ | . | . |
| **G308933A**<br>**MOPB01000004.1** | **R570H** | **BJI72_0342**<br>**grlA** | **DNA topoisomerase**<br>**IV subunit A** | . | √ | √ | √ | √ | √ | √ |
| T13964TTA<br>MOPB01000008.1 | V69delinsIRINQ | BJI72_0448<br>purR | pur operon repressor | . | . | . | . | . | √ | √ |
| G52A<br>MOPB01000047.1 | Synonymous<br>mutation | BJI72_2594<br>sdrE | serine-aspartate<br>repeat-containing<br>protein E | . | √ | √ | √ | √ | √ | . |

Bolded entries represent SNP, variants, and amino acid changes that appeared in all *S. aureus*-1 and 2 strains, but not in *S. aureus* ATCC 29213.
√, the mutation present; ., no mutation.

### Mutations observed in the quinolone-resistance determining regions (QRDRs)

Target sequencing of QRDRs indicated that there were some mutations in both *grlA* (S80F) and *gyrB* (T451S and/or R450S) (***Table 4***), among those, mutations of both genes were found in CIP- and OFL-1 while only one mutation in *gyrB* (T451S) was found in LEV-1. Additional sequencing of the *S. aureus* strains at days 4^th^, 6^th^, 8^th^, 10^th^ of CIP-, LEV- and OFL-exposure revealed the *grlA* mutation (S80F) to appeared in earlier steps than the ones in *gyrB*, suggesting the primary role of *grlA* mutation in FQ resistant phenotype (***Table 4***).

**Table 4.**
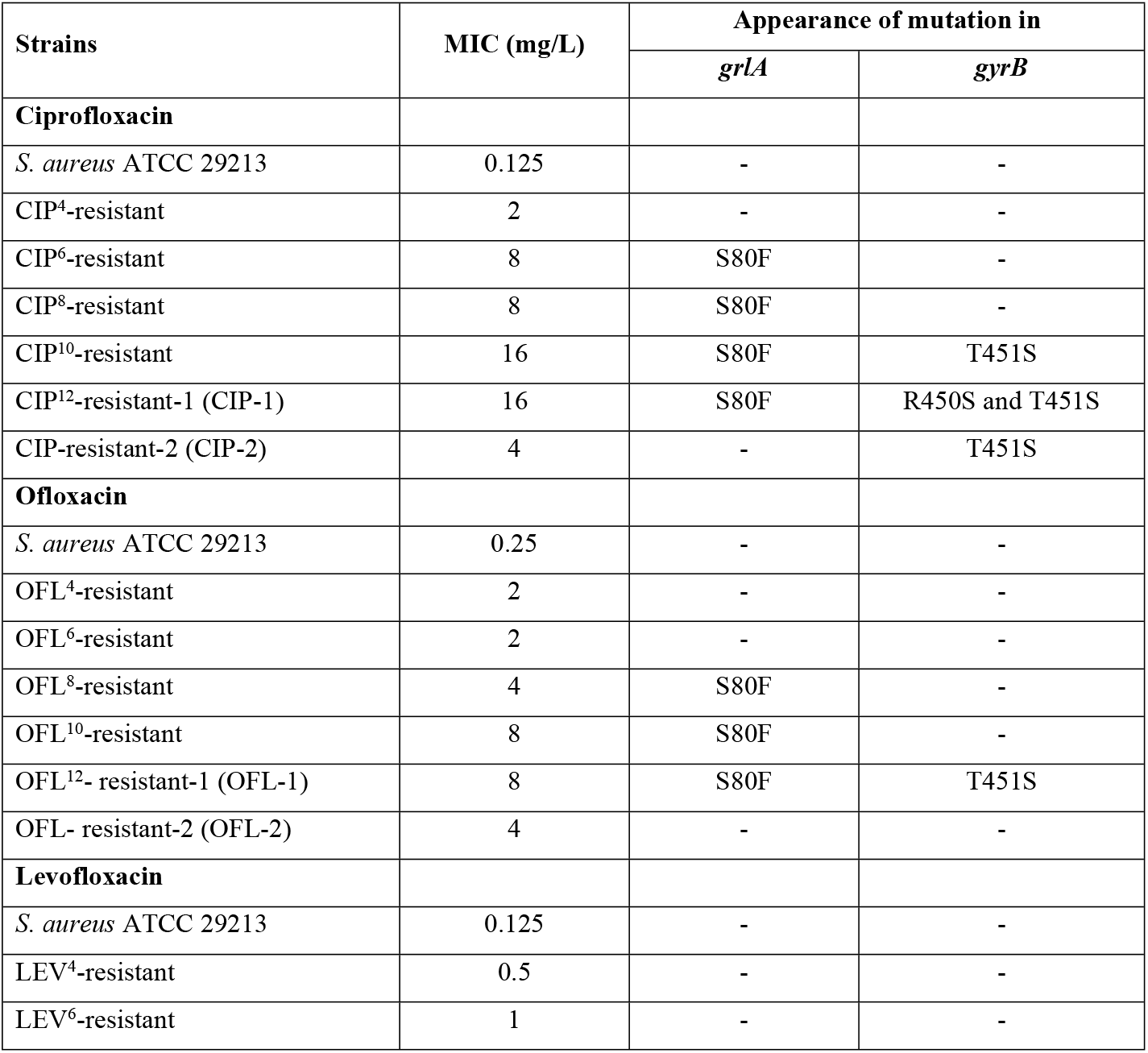
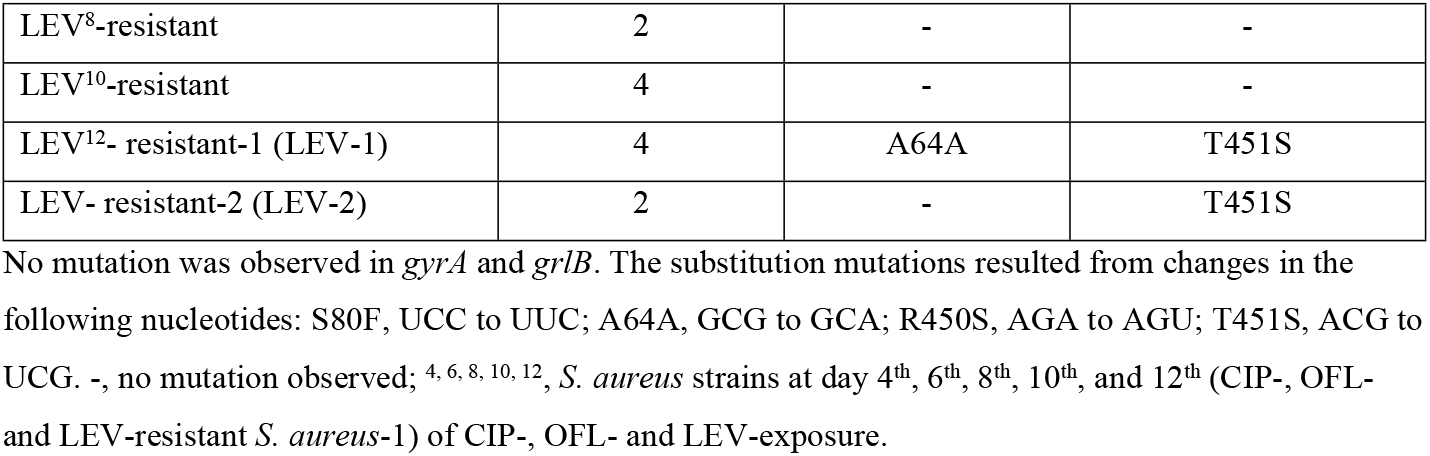
Sequencing result of *grlA* and *gyrB* of *S. aureus* strains.

| Strains | MIC (mg/L) | Appearance of mutation in |  |
| --- | --- | --- | --- |
|  |  | *grlA* | *gyrB* |
| **Ciprofloxacin** |  |  |  |
| *S. aureus* ATCC 29213 | 0.125 | - | - |
| CIP<sup>4</sup>-resistant | 2 | - | - |
| CIP<sup>6</sup>-resistant | 8 | S80F | - |
| CIP<sup>8</sup>-resistant | 8 | S80F | - |
| CIP<sup>10</sup>-resistant | 16 | S80F | T451S |
| CIP<sup>12</sup>-resistant-1 (CIP-1) | 16 | S80F | R450S and T451S |
| CIP-resistant-2 (CIP-2) | 4 | - | T451S |
| **Ofloxacin** |  |  |  |
| *S. aureus* ATCC 29213 | 0.25 | - | - |
| OFL<sup>4</sup>-resistant | 2 | - | - |
| OFL<sup>6</sup>-resistant | 2 | - | - |
| OFL<sup>8</sup>-resistant | 4 | S80F | - |
| OFL<sup>10</sup>-resistant | 8 | S80F | - |
| OFL<sup>12</sup>- resistant-1 (OFL-1) | 8 | S80F | T451S |
| OFL- resistant-2 (OFL-2) | 4 | - | - |
| **Levofloxacin** |  |  |  |
| *S. aureus* ATCC 29213 | 0.125 | - | - |
| LEV<sup>4</sup>-resistant | 0.5 | - | - |
| LEV<sup>6</sup>-resistant | 1 | - | - |
| LEV<sup>8</sup>-resistant | 2 | - | - |
| LEV<sup>10</sup>-resistant | 4 | - | - |
| LEV<sup>12</sup>- resistant-1 (LEV-1) | 4 | A64A | T451S |
| LEV- resistant-2 (LEV-2) | 2 | - | T451S |

No mutation was observed in *gyrA* and *griB*. The substitution mutations resulted from changes in the following nucleotides: S80F, UCC to UUC; A64A, GCG to GCA; R450S, AGA to AGU; T451S, ACG to UCG. -, no mutation observed; <sup>4, 6, 8, 10, 12</sup>, *S. aureus* strains at day 4<sup>th</sup>, 6<sup>th</sup>, 8<sup>th</sup>, 10<sup>th</sup>, and 12<sup>th</sup> (CIP-, OFL- and LEV-resistant *S. aureus*-1) of CIP-, OFL- and LEV-exposure.

### Overexpression of alternative sigma factors in FQ-exposed strains

The data in ***Fig 2*** illustrates that there were upregulations of *sigB* and *sigS* genes in most FQ-exposed strains, except for OFL-2 and LEV-2. The expression of *sigB* and *sigS* genes were recorded highest in LEV-1, at a mean fold change expression of 2.39 and 3.80 respectively, while the most significant downregulation of both genes was recorded in OFL-2 which were about 0.41 and 0.55 fold decrease. In terms of the *S. aureus*-2 group, the upregulation of both genes in CIP-2 made this the only strain in the group that experienced a rise in gene expression of both alternative sigma factors, σ^B^ and σ^S^ in comparison with initial *S. aureus*. In the statistical analysis between two groups of *S. aureus*-1 and *S. aureus*-2 for the same antibiotic, it can be seen that there was a decrease in gene expression in both *sigB* and *sigS* genes of *S. aureus*-2 strains compared with *S. aureus*-1 strains.

**Fig 2.**
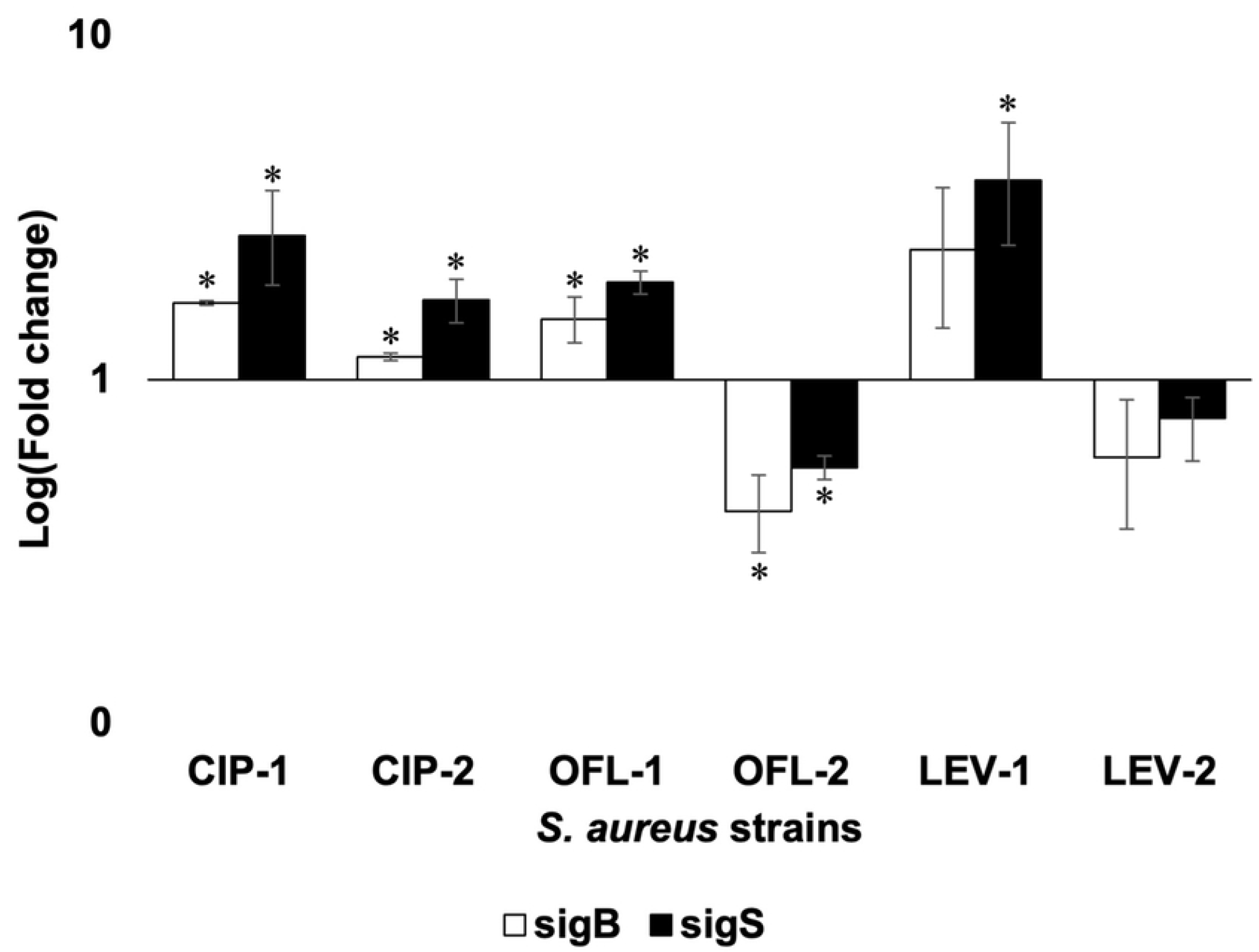
RT-qPCR analysis of *sig B* and *sigS* in *S. aureus* ATCC 29213 and its exposed strains. Fold change was calculated against initial *S. aureus* and visualized on a log scale, with gene expression of initial *S. aureus* as 1. Values greater than one were up-regulated, while values smaller than 1 were down-regulated. Foldchange and confidence level 95% CI (error bar) were calculated in MS Excel according to the standard practice [24]. * indicated a significant difference in gene expression between initial *S. aureus* and FQ-exposed *S. aureus* strains.

### Efflux expression increased in FQ-exposed strains

The effect of FQ exposure on the expression level of the efflux pump genes was checked and compared to initial *S. aureus*. As shown in ***Fig 3***, all *S. aureus*-1 and 2 strains increased the expression of *mgrA, norA, norB*, and *norC*. Regarding to the expression level of *mgrA*, which is involved in the efflux pump regulation of *S. aureus*, LEV-1 exhibited the highest expression level (about 12.63 folds change), followed by CIP-1 and LEV-2 (**Fig 3*A***). Besides, *norA* was also upregulated in all FQ-exposed strains, especially in CIP-1 and CIP-2 of which the expression increased by about 57.61 and 70.78 folds respectively compared to initial strains (**Fig 3*B***). Among *S. aureus*-1 strains, OFL-1 showed the lowest expression in *mgrA, norA, norB*, and *norC*. A similar result was also found in OFL-2 when compared with the remaining *S. aureus*-2 strains. In addition, almost all strains belonging to *S. aureus*-1 group had higher expression in *mgrA, norA, norB*, and *norC* compared to the strains which were treated with the same antibiotic in *S. aureus*-2 group.

**Fig 3.**
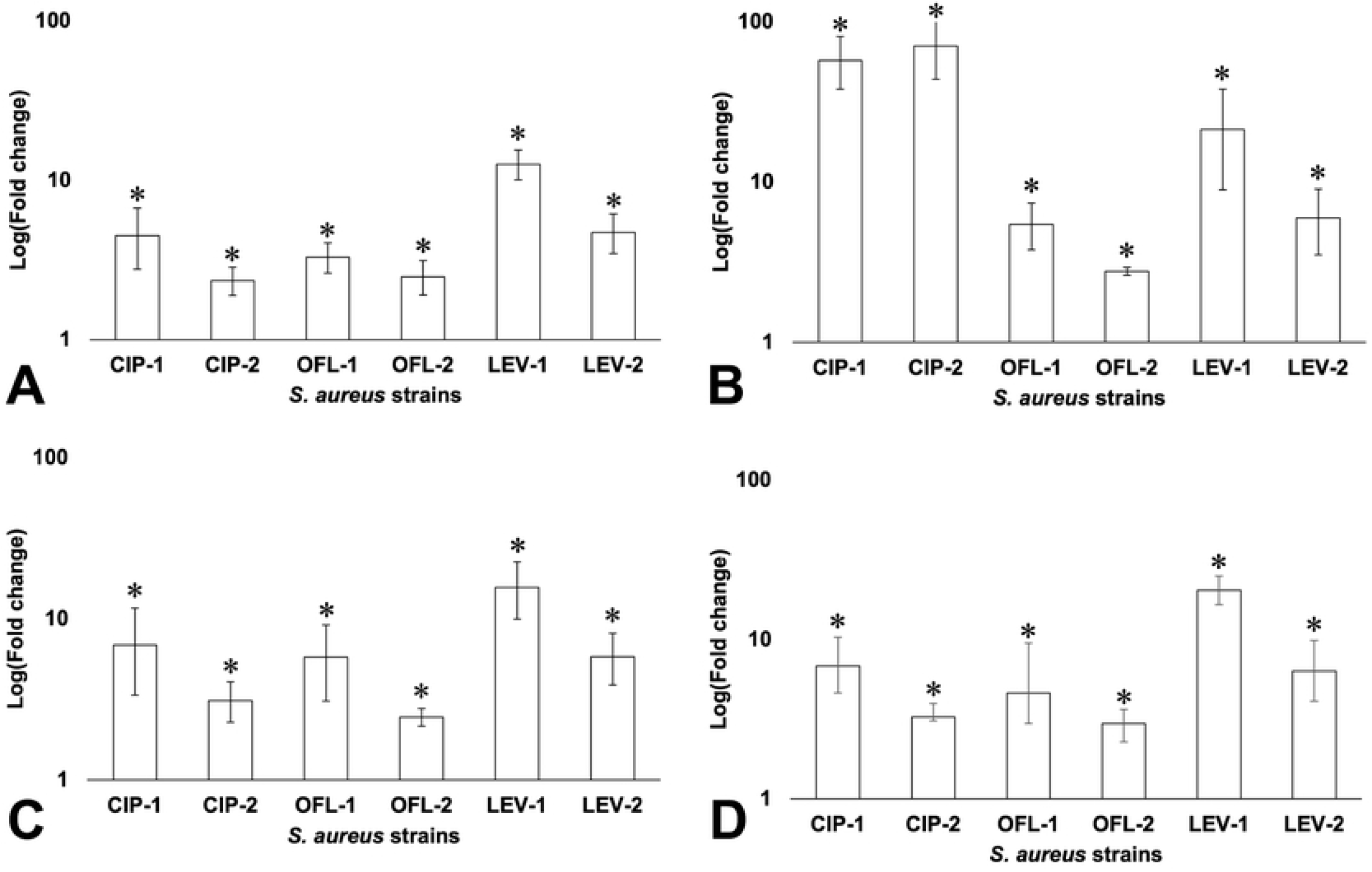
RT-qPCR analysis of *mgrA* (A) and multidrug efflux pump *norA* (B), *norB* (C), and *norC* (D) in *S. aureus* ATCC 29213 and its exposed strains. Fold change was calculated against initial *S. aureus* and visualized on a log scale, with gene expression of initial *S. aureus* as 1. Values greater than one were up-regulated, while values smaller than 1 were down-regulated. Foldchange and confidence level 95% CI (error bar) were calculated in MS Excel according to the standard practice [24]. * indicated a significant difference in gene expression between initial *S. aureus* and FQ-exposed *S. aureus* strains.

### Overexpression of genes related to FQ-resistant development in *S. aureus*

According to the WGS results, there were genomic mutations located in the sequence of *rimI* and *fmtB* in both *S. aureus*-1 and -2 strains. Besides, 7 SNPs were also found in all FQ-exposed strains which were located in the upstream region of *norA* and the gene encoding hypothetical protein BJI72_0645 (***Table 3***). The results suggested that these variants might be related to FQ-resistant development in *S. aureus*. Therefore, investigating the expression of *rimI, fmtB*, and the gene encoding hypothetical protein BJI72_0645 should be carried out to better understand the relationship between these mutations and the FQ-resistant development in *S. aureus*.

Under the effects of FQ exposure, *rimI* was overexpressed in all FQ-exposed strains, especially the expression increased by 30.43 folds in LEV-1 (***Fig 4A***). The data also indicated that *fmtB* was upregulated in *fmtB*-mutant strains (***Fig 4B***). Besides, although there was no *fmtB* mutation in CIP-1 and CIP-2 which were resistant to ciprofloxacin, *fmt*B was also overexpressed in both strains. Regarding the expression level of the gene encoding hypothetical protein BJI72_0645, this gene was also overexpressed in all FQ-exposed strains, especially LEV-1 (***Fig 4C***). In addition, this study also found that the expression of *rimI, fmtB*, and the gene encoding hypothetical protein BJI72_0645 in *S. aureus*-1 strains was higher than that in corresponding *S. aureus*-2 strains in all cases.

**Fig 4.**
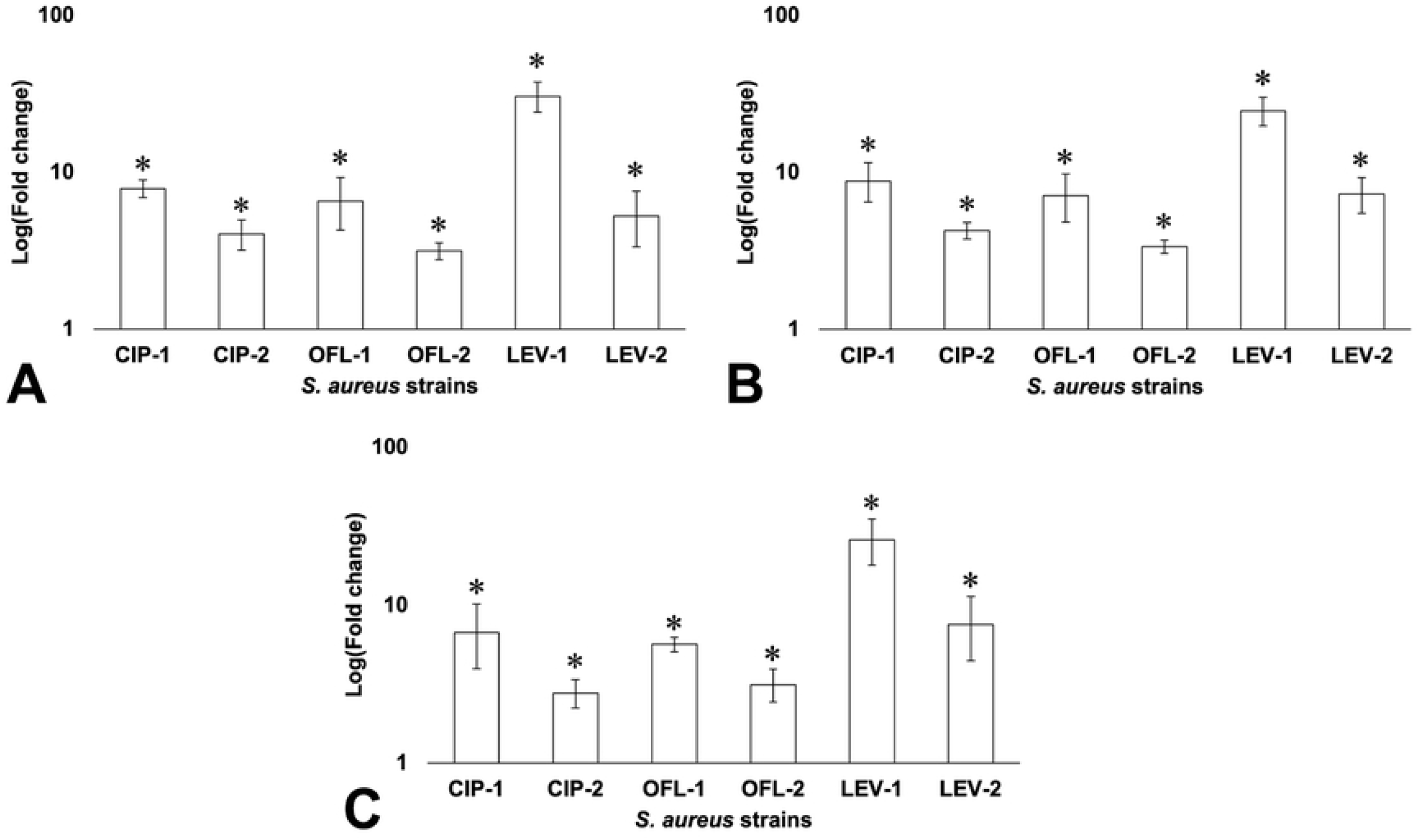
RT-qPCR analysis of *rimI* (A), *fmtB* (B), and the gene encoding hypothetical protein BJI72_0645 (C) in *S. aureus* ATCC 29213 and its exposed strains. Fold change was calculated against initial *S. aureus* and visualized on a log scale, with gene expression of initial *S. aureus* as 1. Values greater than one were up-regulated, while values smaller than 1 were down-regulated. Foldchange and confidence level 95% CI (error bar) were calculated in MS Excel according to the standard practice [24]. * indicated a significant difference in gene expression between initial *S. aureus* and FQ-exposed *S. aureus* strains.

## Discussion

### Sub-MIC exposure to FQ altered the antibiotic susceptibility profile of *S. aureus*

After 12 days of exposure to the sub-MICs of FQs, *S. aureus* increased its MIC values and became resistant to exposed antibiotics. Serial exposure to sub-MICs can create positive selection pressure that drove the development of resistant phenotypes, which is in agreement with previous literature [25–27]. Although *S. aureus* had a decrease in MIC value after 10 days of culturing in an antibiotic-free medium, indicating some reversion of resistance trait, *S. aureus*-2 strains kept their resistance to FQs. Besides, after being serial exposed to ciprofloxacin, levofloxacin, and ofloxacin, *S. aureus* became resistant not only to the exposed antibiotic and its related FQs but also to other antibiotics of unrelated groups such as ampicillin, amoxicillin, doxycycline, erythromycin, and lincomycin.

### Genomic alterations involved in FQ-resistant development in *S. aureus*

It has been shown that the acquisition of FQ resistance is mainly due to the mutations in target enzymes, topoisomerase IV (GrlA/B) as well as DNA gyrase (GyrA/B) [8]. In *S. aureus*, the mutations occurred more frequently in QRDRs of *grlA* (S80F or Y, E84K, and A116E or P) and *gyrA* (S84L or A, S85P, and E88K) which were described as the primary FQ resistance mechanism [8]. Other studies have found that the mutations in QRDRs of *gyrB* including D437N, R458E, D432N and N470D also contributed to FQ resistance [11,12].

In our *in vitro* selected FQ-resistance model, we found mutations in both *grlA* (S80F) and *gyrB* (T451S and/or R450S), among those, mutations of both genes were found in CIP- and OFL-1 while only one mutation in *gyrB* (T451S) was found in LEV-1. Additional sequencing of the *S. aureus* strains at day 4^th^, 6^th^, 8^th^, 10^th^ of CIP-, LEV- and OFL-exposure revealed the *grlA* mutation (S80F) to be appeared in earlier steps than the ones in *gyrB*, suggesting the primary role of *grlA* mutation in FQ resistant phenotype. These findings suggested that the mutation in *grlA* (S80F) primarily induced the emergence of FQ resistance and might support the mutations in *gyrB* (T451S); which was required for high-level FQ resistance in *S. aureus*.

In our study, we observed the resistance level for the occurrence of *grlA* mutation only and both *grlA* and *gyrB* seemed to be similar to the reported information [4,28]. Interestingly, we observed the revertance of all mutations in *grlA* (turning back to the wild-type genotype) without a significant MIC decrease in OFL- and LEV-2 and only a four-time reduction in CIP-revertant strain when both mutations in *grlA* (S80F) and in *gyrB* (R450S) were together disappeared. Moreover, even though OFL-2 kept most of its resistant ability, no mutation in *gyrA, gyrB, grlA*, and *grlB* QRDRs was retained. Our results were similar to the observation in *E. faecium* of Yoshihiro *et al*. who proposed that other resistance mechanism(s) such as multidrug resistance efflux pumps might be involved in the development of FQ resistance [29].

### Changes in expression of alternative sigma factors associated with the adaptive response of *S. aureus* under serial FQ exposure

There is a long-established correlation between increased resistance and fitness cost [30]. If the FQ-resistance mechanism imposes a fitness cost, it would be advantageous for bacteria to be able to exhibit the resistance phenotype only when the threat exists. Even if there is no fitness cost, the production of resistance-mediating proteins could impose an unnecessary metabolic burden on the bacteria in an antimicrobial-free environment. From the transcriptional analysis of *sigB* and *sigS* genes, we can see that both genes were downregulated or reduced in gene expression in the *S. aureus-2* group compared with the *S. aureus*-1 group. The antibiotic-free media removed the FQ stressors that induced the adaptive response, which in turn deprived the need for upregulation of *sigB* and *sigS* genes. This suggests that both *sigB* and *sigS* also present a selective advantage to *S. aureus* and play an important role in *S. aureus* fitness and adaptation to stress.

### Sub-MIC exposure to FQ led to overexpression of the efflux pump and its regulator

Several specific efflux pumps have been associated with antibiotic resistance in clinical isolates of *S. aureus*. With respect to FQ, *norA, norB*, and *norC* are the most important efflux pumps localized on the cytoplasmic membrane of *S. aureus* [21]. These pumps can extrude many chemically and structurally dissimilar compounds, namely hydrophilic and hydrophilic fluoroquinolones (such as norfloxacin, ciprofloxacin, moxifloxacin, and garenoxacin), dyes (like ethidium bromide and rhodamine) and biocides (tetraphenylphosphonium and cetrimide) [31,32]. It has been shown that *mgrA* functions as a positive regulator of *norB* and a negative regulator of *norA* and *norC* [33].

Regarding the expression level of *mgrA*, the results in this study were consistent with our previous study on proteomic analysis [34]. In order to test the effects of overexpression of *mgrA* on multidrug efflux pump *norA, norB*, and *norC*, RT-qPCR was applied for determining their expression. In analysis, the expression of norA, *norB*, and *norC* was increased in all tested strains. It means that efflux pumps were activated and could be one of the mechanisms leading to the occurrence of multidrug-resistant *S. aureus* strains. Our results are in agreement with the previous study, in which overexpression of MgrA has led to a change in the expression level of *norA* and *norB* [32]. However, the increased expression of *norC* in this study was not associated with the regulation of *mgrA*, and it might result from mutational alterations in uncharacterized loci that affect the expression of this gene.

The decrease in efflux expression in *S. aureus*-2 compared to those of *S. aureus*-1 suggests that under the FQ stressor in the environment, the efflux activity of *S. aureus* was promoted as an adaptive response to resist the antibiotics. However, the efflux activity could decrease when *S. aureus*-1 was cultured continuously in antibiotic-free media even though it still kept its resistance to FQ. This is consistent with the result that *S. aureus* turned to an FQ-resistant phenotype after 12-day serial exposure to FQ, and the MIC values witnessed a slight decrease after 10-day continuous culturing *S. aureus*-1 in antibiotic-free media.

### Sub-MIC exposure to FQs affected the protein acetylation and multi-drug resistant phenotype of *S. aureus*

Protein acetylation is one of the major post-translational modifications (PTMs) which play an important role in cell signaling and occur when the cell encounters specific environmental stress conditions [35–37]. RimI encoded by *rimI* is a ribosomal protein S18 acetyltransferase which catalyzes the acetylation of the N-terminal alanine of ribosomal protein S18; and also acts as an N-epsilon-lysine acetyltransferase that catalyzes acetylation of several proteins [38,39]. In this study, an insertion mutation (S33_S34insC) in *rimI* was found in all *S. aureus*-1 and 2 that might affect the RimI activities and the protein acetylation in *S. aureus*. In terms of expression analysis, *rimI* was overexpressed in these mutant strains compared to initial *S. aureus*. These results suggested that the mutation in *rimI* as well as its overexpression in FQ-exposed *S. aureus* was an adaptive response under antibiotic stressors. This might be associated with acetylation which alters mRNA translation efficiency in several proteins in *S. aureus* to maintain its survival in an antibiotic-stress environment.

The *fmtB* gene codes for FmtB, a ∼263 kDa cell wall-anchored protein [40]. Although there is limited information about the function of *fmtB*, it has been proven to contribute to the methicillin-resistant phenotype in *S. aureus* [41,42]. According to the antibiotic susceptibility profile of seven *S. aureus* strains, after being exposed to FQ, *S. aureus* enhanced its resistance to FQ and other antibiotics including B-lactam. Besides, the mutation K1283delinsRG (CTT26104C) in *fmtB* sequence was also found in OFL-and LEV-exposed strains. For RT-qPCR analysis, *fmtB* was upregulated in all mutant strains. Although there was no mutation on *fmtB* found in CIP-1 and CIP-2, which were resistant to ciprofloxacin, *fmtB* was also overexpressed in both strains. It was suggested that the *in vitro*-induced FQ resistance cooperating with the acetylation modification somehow affects the *fmtB* expression and the antibiotic resistance of *S. aureus* to FQs and other antibiotics of unrelated groups leading to a multi-drug resistance phenotype.

According to whole genome analysis, 7 SNPs were found in the upstream region of both *norA* and the gene encoding hypothetical protein BJI72_0645. The RT-qPCR results indicated that the variants might play a crucial role in regulating the transcription of these genes because both were overexpressed in all FQ-exposed *S. aureus*.

### From genetic alterations to protein expression

The development of antibiotic-resistant might involve not only transcriptional regulation and genetic modifications but also translational regulation. In the proteomic study of our research group, under the FQ stressor in the environment, there were 147 unique proteins in *S. aureus*-1 and *S. aureus*-2 which changes their expression in comparison to initial strain [34]. Regarding the molecular function of differently expressed proteins, 93 proteins were responsible for binding, 89 proteins for catalytic activity, 28 proteins for the structural constituent of ribosome, and 10 other proteins were involved in transcription factor activity (6), transporter activity (2) and antioxidant activity (2). Proteins involved in SOS/stress response and antibiotic resistance and pathogenesis were upregulated upon FQ exposure. Among them, RecA and MgrA are global regulators which are strongly implicated in antibiotic resistance development. The presence of RecA leads to the upregulation of SOS genes which involve in DNA repair and promote the autoproteolysis cleavage of LexA (an SOS gene repressor) [43,44]; while MgrA affects multiple *S. aureus* genes which encoded proteins involving multidrug resistance, autolytic activity, and virulence [45,46]. The fluctuation in protein expression depends on many factors including extracellular agents such as harsh temperatures, chemical and antibiotic stress in the environment, and intracellular agents such as modifications in the genetic materials. Previous studies have proven that the elements belonging to genetic code including amino acid [47], untranslated regions [48–51], length [51], GC content [52–54], and mRNA secondary structure [55] affected the regulation of protein expression. However, under the serial exposure of *S. aureus* to FQs, the direct relationship between protein expression and genetic modifications found in this study was not clear. The results suggest that resistance to multiple drugs including FQs under drug exposure generally requires the contribution of multiple proteins and processes, thus the genetic mutations found in FQ-exposed *S. aureus* might indirectly affect the expression of resistance-associated proteins that made these proteins more active to help this bacterium adapt the antibiotic-containing environments.

### Conclusion

Exposure to sub-MICs of FQs provided positive selection pressure for the development of resistance traits leading to alterations to the antibiotic susceptibility, growth rate, and morphology of the pathogen. These alterations were reflected in the DNA mutations and the change at mRNA expression levels. Multidrug resistance or antibiotic-resistant development is a complicated and multifactorial process. The found mutations might be spontaneous, but their combination is clearly a cause of the resistance status observed in the bacterium.

## Supporting information

**S1 Table.**
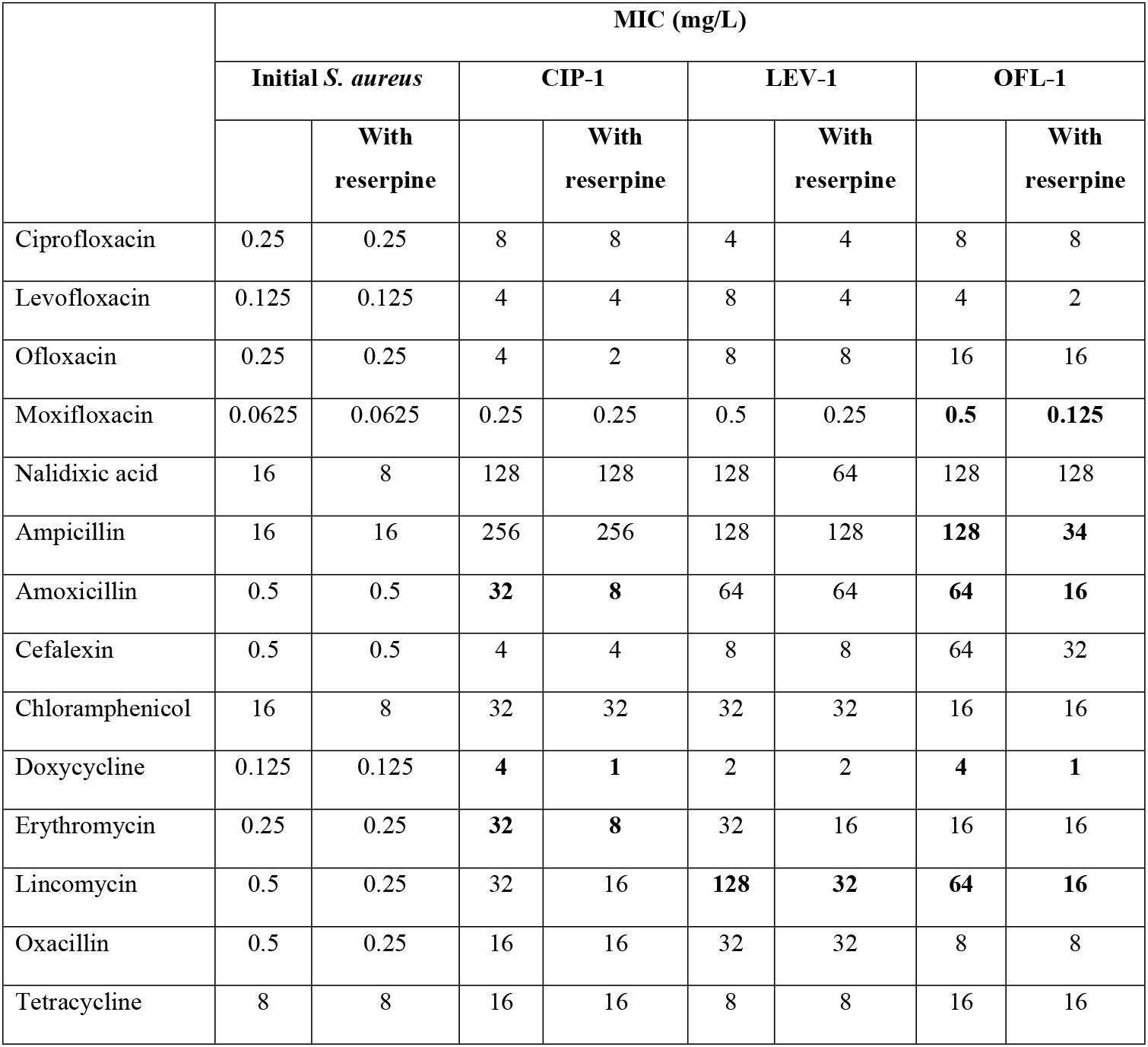
The effect of reserpine on antibiotic susceptibility profile of initial and *S. aureus*-1 strains.

**S2 Table.**
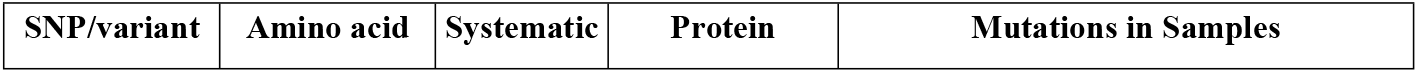
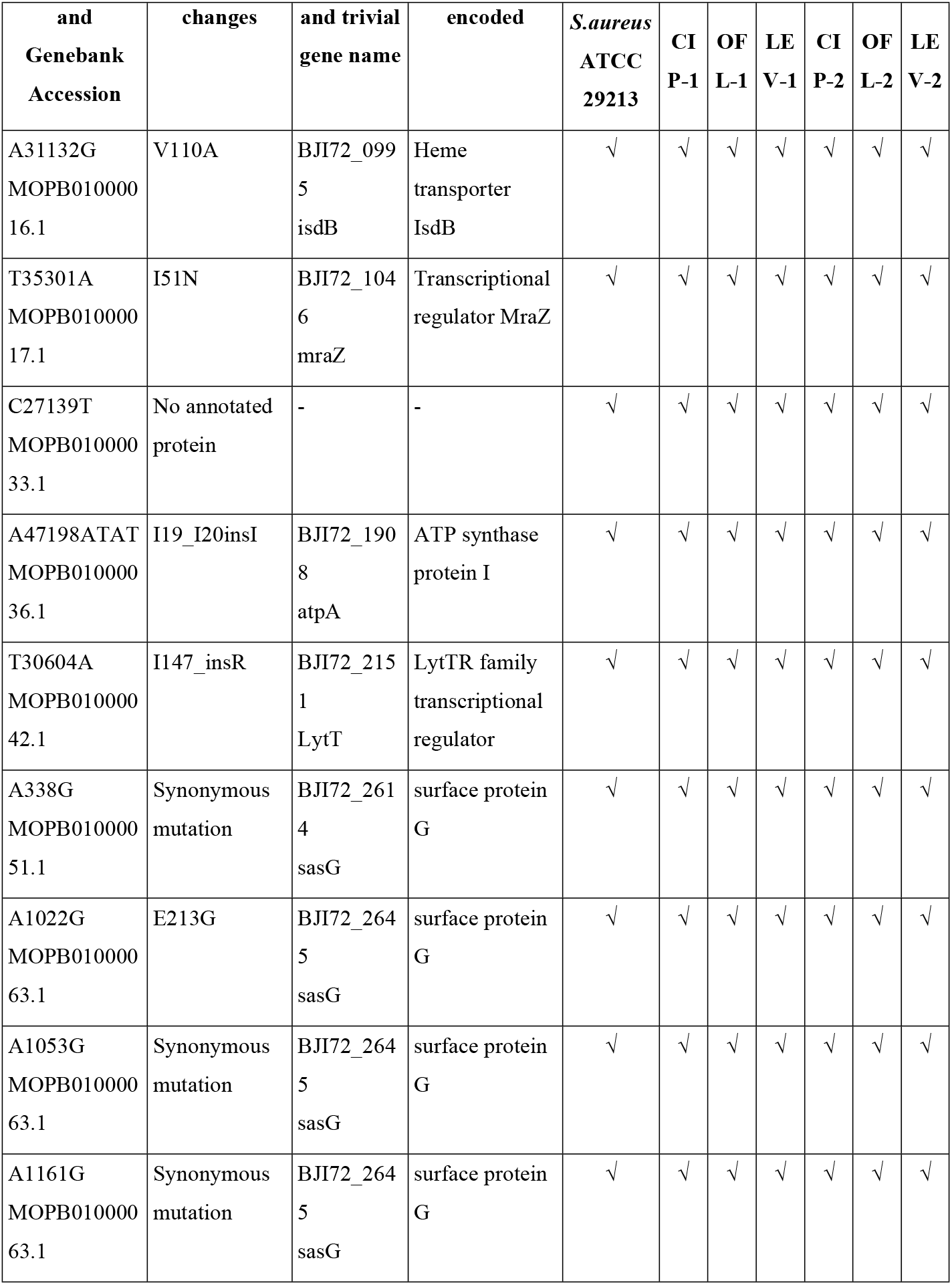
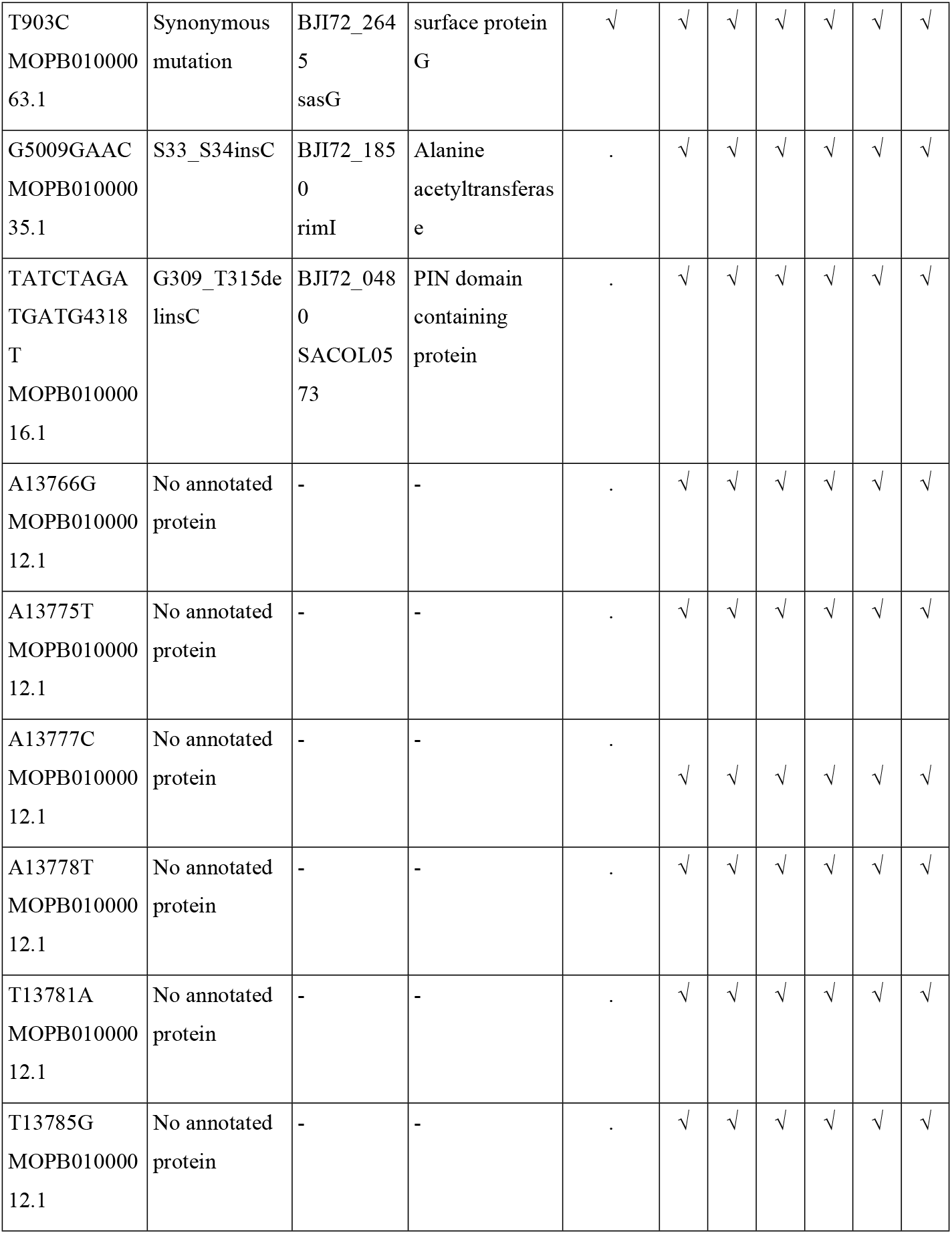
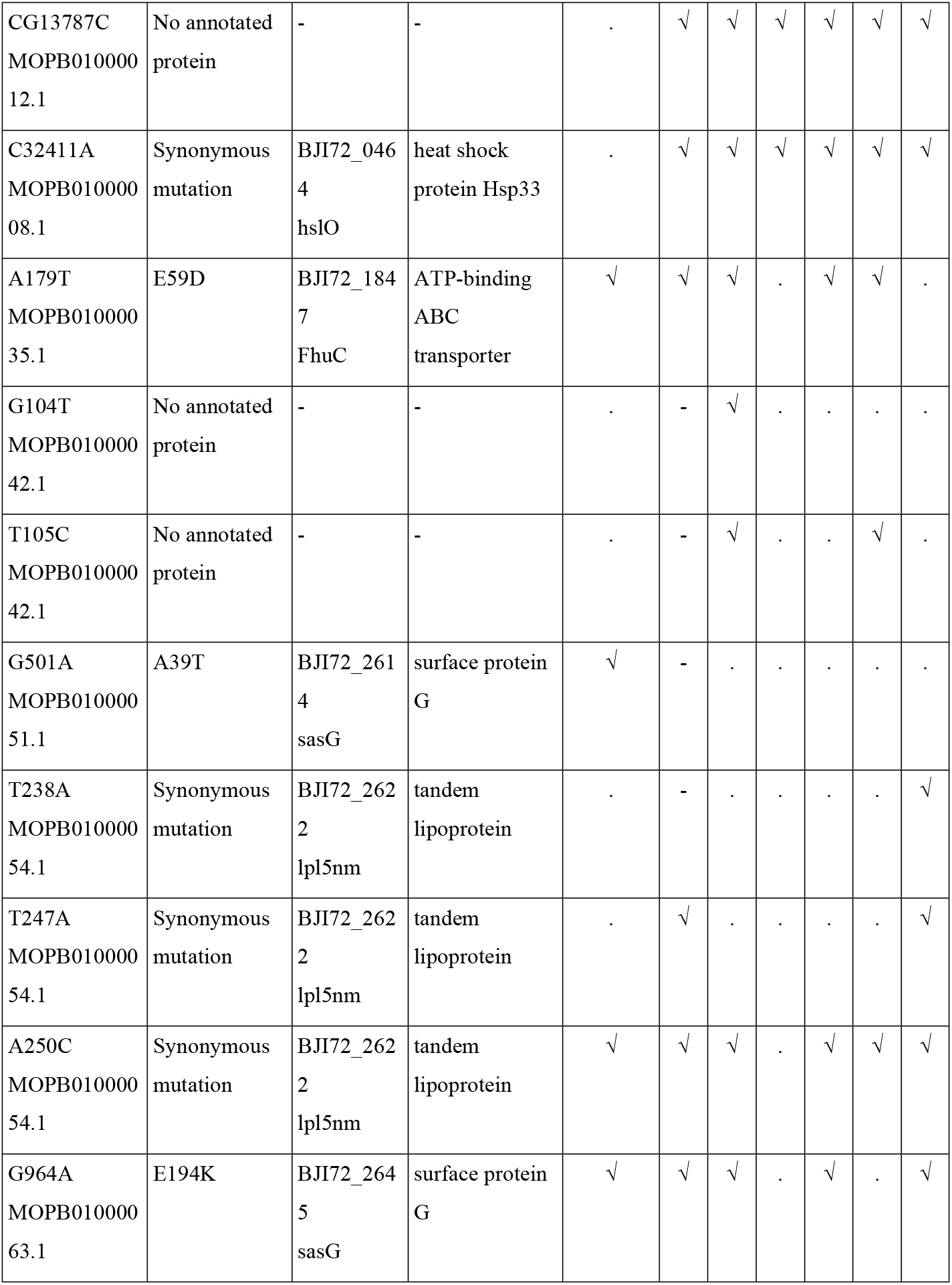
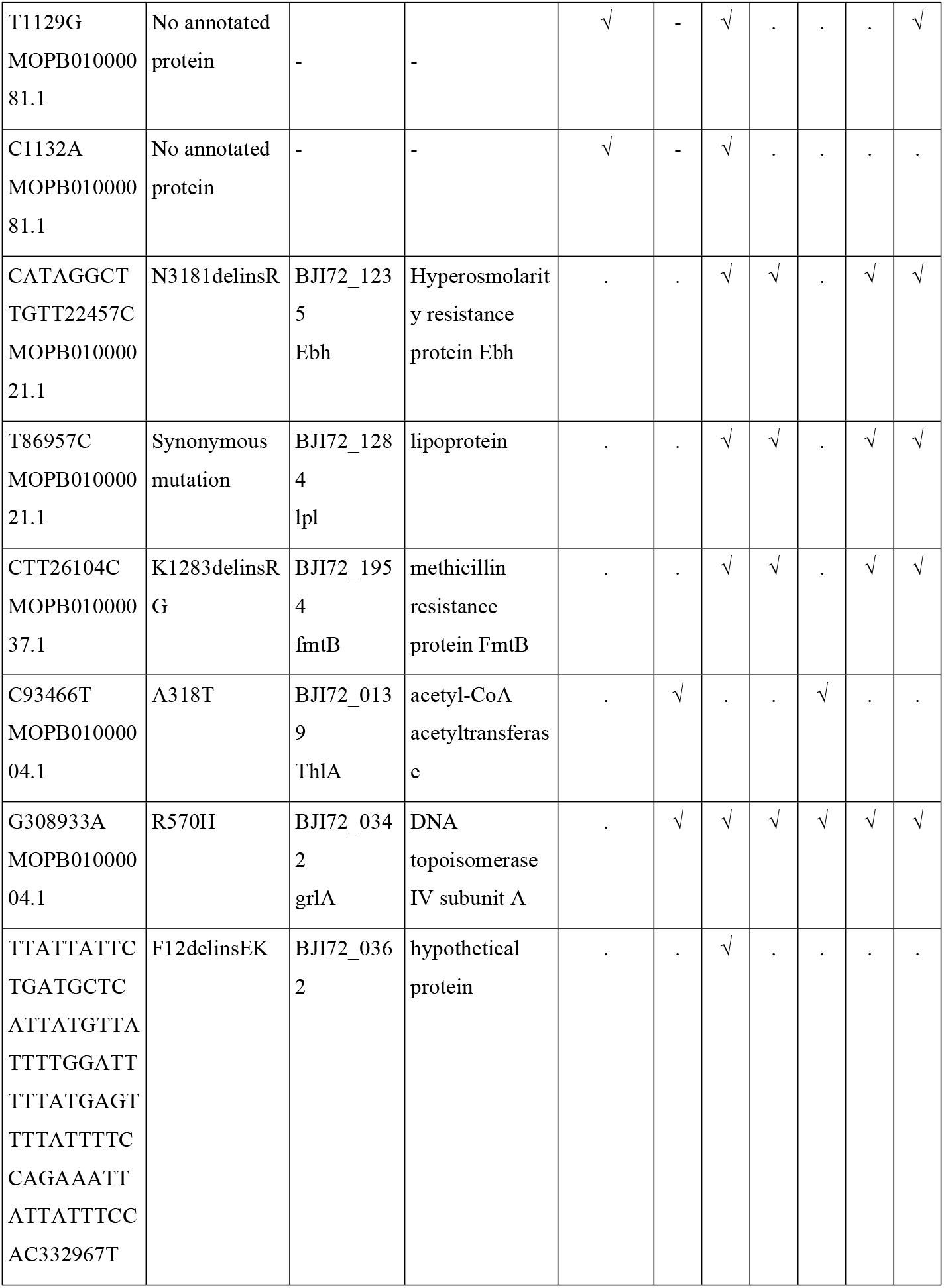
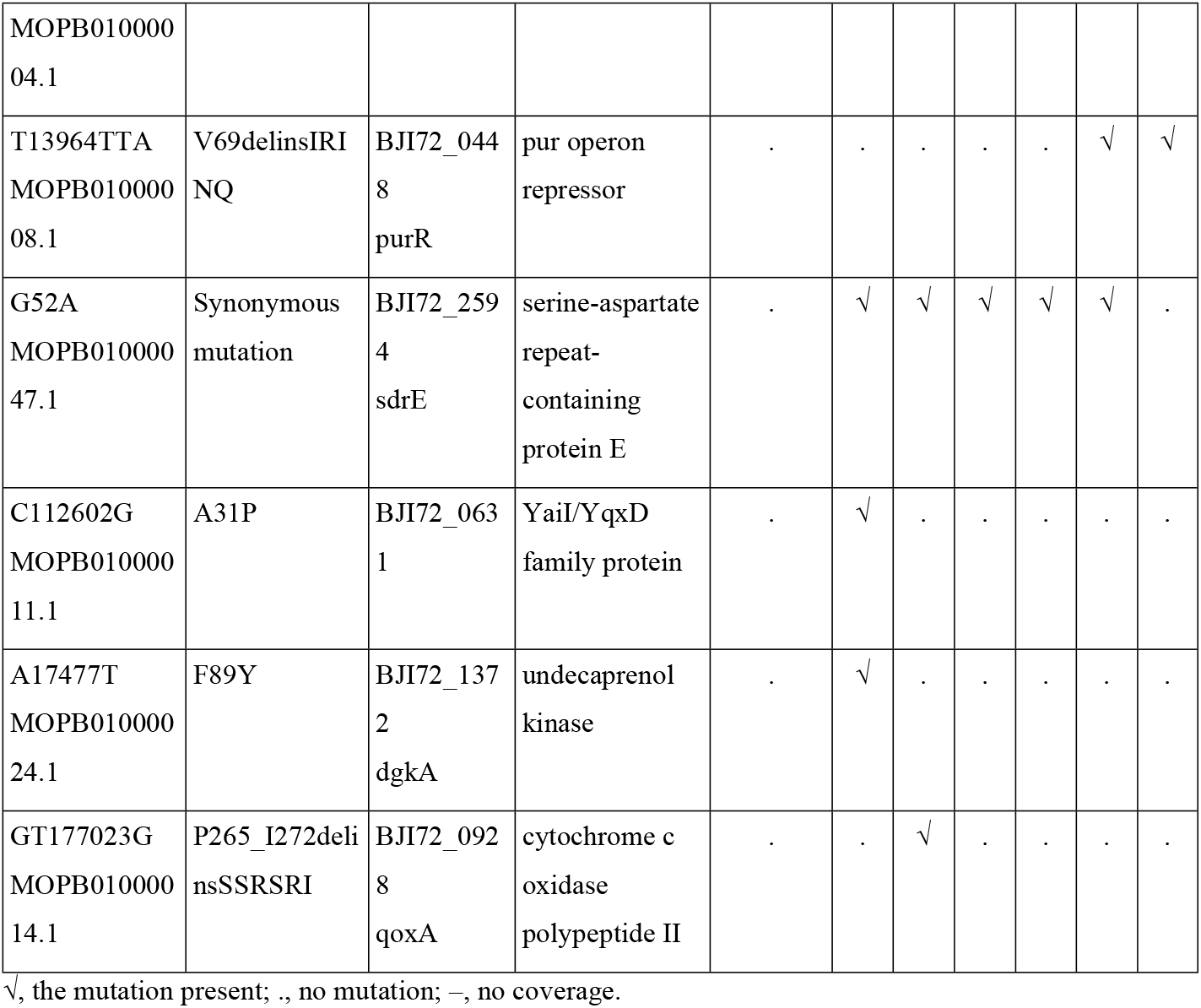
List of single nucleotide polymorphisms (SNPs), variants, and amino acid changes found in *S. aureus*-1 (CIP-1, OFL-1, and LEV-1) and *S. aureus*-2 (CIP-2, OFL-2, and LEV-2) strains but not in *S. aureus* ATCC 29213.

**S3 Table.**
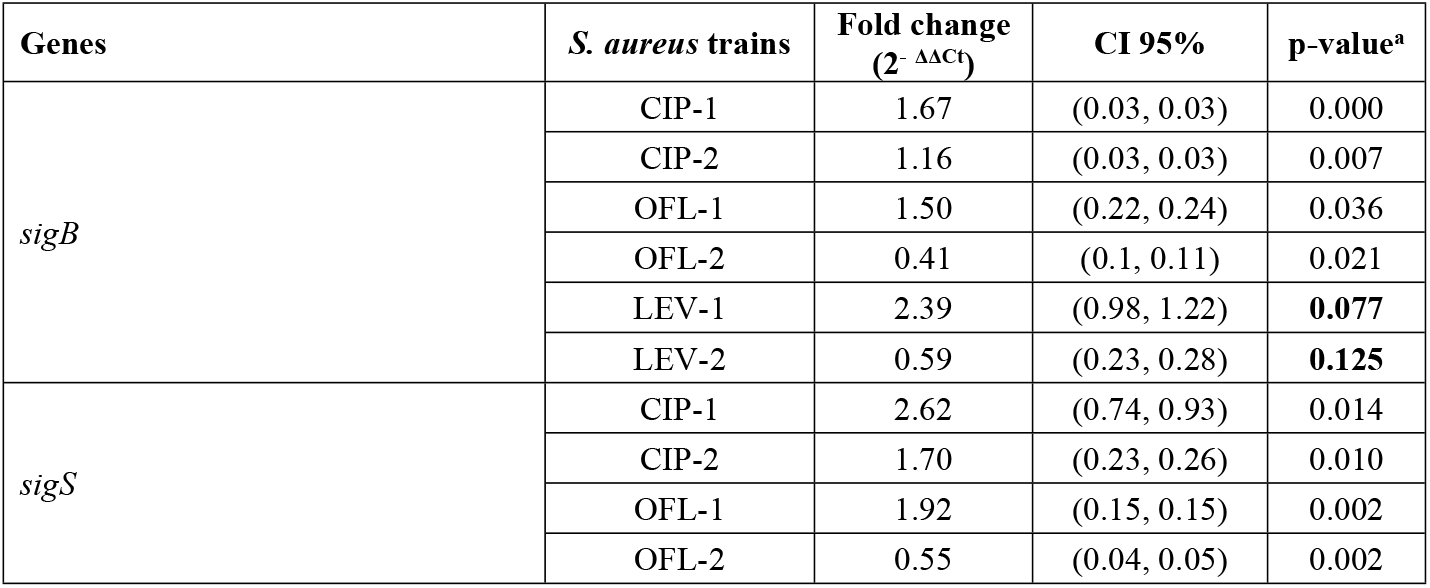
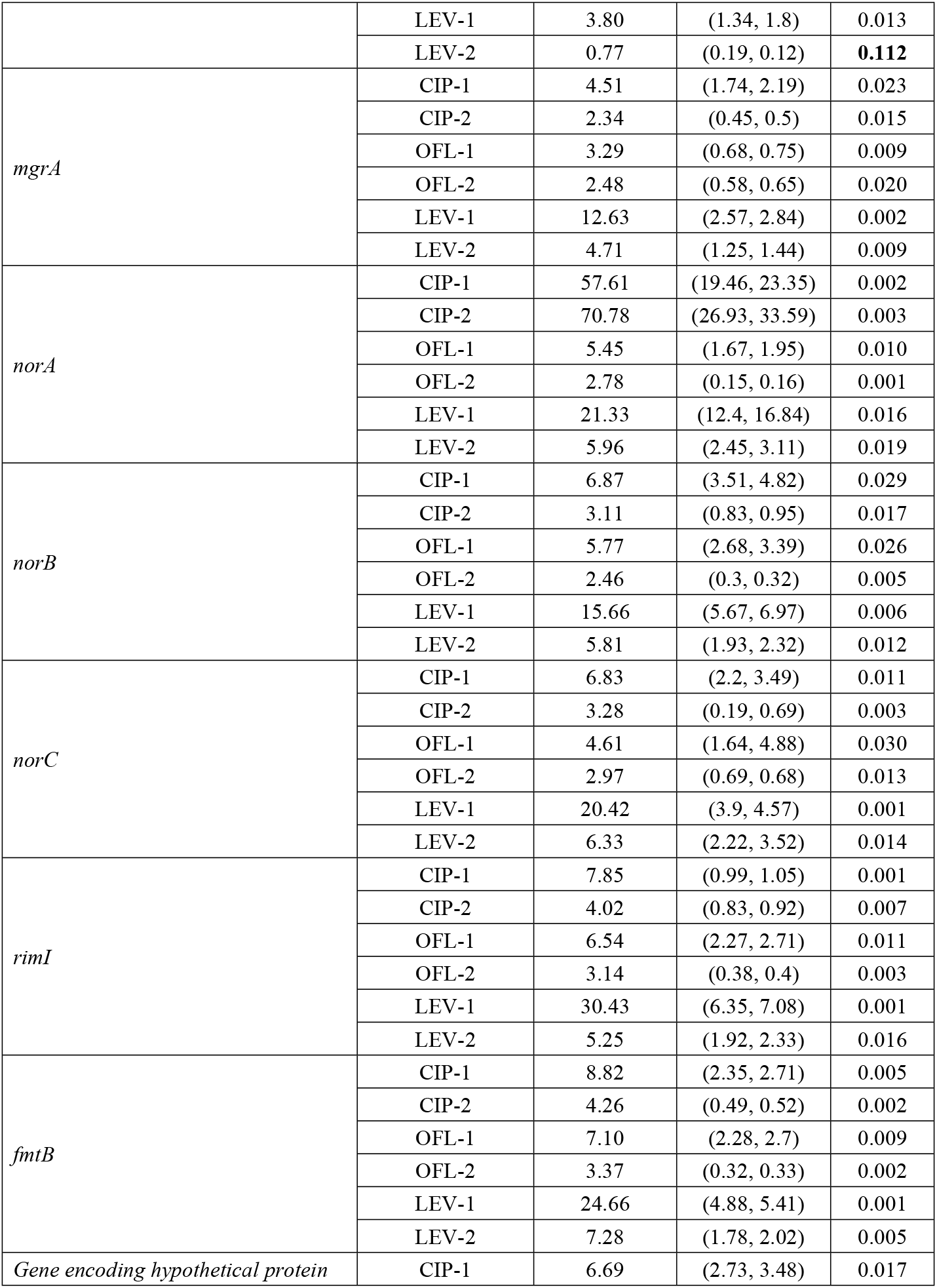
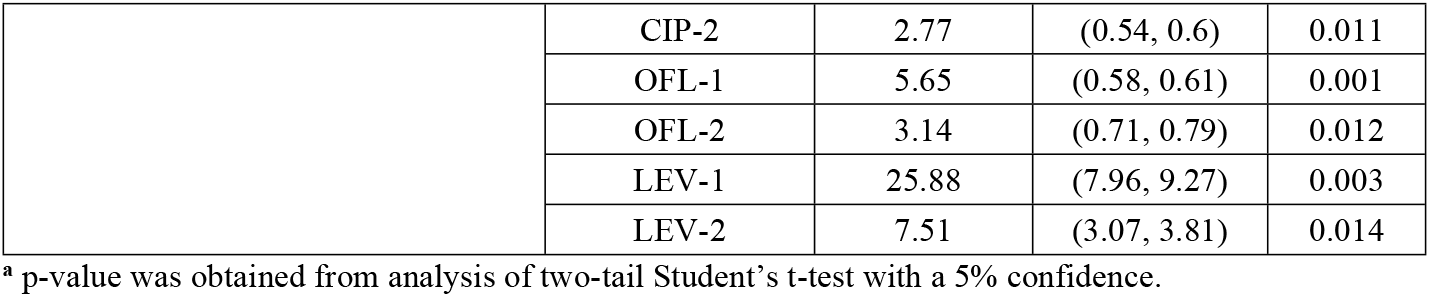
**The statistical analysis of fold change in gene expression between initial *S. aureus* strain and FQ-exposed *S. aureus* strains** (mean ± standard error).

## References

[1] Gordon RJ, Lowy FD. Pathogenesis of Methicillin-Resistant Staphylococcus aureus Infection. Clin Infect Dis 2008;46:S350. https://doi.org/10.1086/533591.

[2] Travers K, Barza M. Morbidity of infections caused by antimicrobial-resistant bacteria. Clin Infect Dis 2002;34 Suppl 3:S131–4. https://doi.org/10.1086/340251.

[3] Vien LTM, Minh NNQ, Thuong TC, Khuong HD, Nga TVT, Thompson C, et al. The co-selection of fluoroquinolone resistance genes in the gut flora of Vietnamese children. PLoS One 2012;7. https://doi.org/10.1371/JOURNAL.PONE.0042919.

[4] Redgrave LS, Sutton SB, Webber MA, Piddock LJV. Fluoroquinolone resistance: mechanisms, impact on bacteria, and role in evolutionary success. Trends Microbiol 2014;22:438–45. https://doi.org/10.1016/J.TIM.2014.04.007.

[5] Jacoby GA. Mechanisms of resistance to quinolones. Clin Infect Dis 2005;41 Suppl 2. https://doi.org/10.1086/428052.

[6] Hooper DC. Fluoroquinolone resistance among Gram-positive cocci. Lancet Infectious Diseases 2002;2:530–8. https://doi.org/10.1016/S1473-3099(02)00369-9.

[7] Ferrero L, Cameron B, Crouzet J. Analysis of gyrA and grlA mutations in stepwise-selected ciprofloxacin-resistant mutants of Staphylococcus aureus. Antimicrob Agents Chemother 1995;39:1554–8. https://doi.org/10.1128/AAC.39.7.1554.

[8] Kaatz GW, Seo SM. Mechanisms of fluoroquinolone resistance in genetically related strains of Staphylococcus aureus. Antimicrob Agents Chemother 1997;41:2733–7. https://doi.org/10.1128/AAC.41.12.2733.

[9] Piddock LJV. Mechanism of quinolone uptake into bacterial cells. J Antimicrob Chemother 1991;27:399–403. https://doi.org/10.1093/JAC/27.4.399.

[10] Aldred KJ, Kerns RJ, Osheroff N. Mechanism of Quinolone Action and Resistance. Biochemistry 2014;53:1565. https://doi.org/10.1021/BI5000564.

[11] Ito H, Yoshida H, Bogaki-Shonai M, Niga T, Hattori H, Nakamura S. Quinolone resistance mutations in the DNA gyrase gyrA and gyrB genes of Staphylococcus aureus. Antimicrob Agents Chemother 1994;38:2014. https://doi.org/10.1128/AAC.38.9.2014.

[12] Tanaka M, Onodera Y, Uchida Y, Sato K. Quinolone resistance mutations in the GrlB protein of Staphylococcus aureus. Antimicrob Agents Chemother 1998;42:3044–6. https://doi.org/10.1128/AAC.42.11.3044.

[13] Costa S, Falcão C, Viveiros M, MacHado D, Martins M, Melo-Cristino J, et al. Exploring the contribution of efflux on the resistance to fluoroquinolones in clinical isolates of Staphylococcus aureus. BMC Microbiol 2011;11:1–12. https://doi.org/10.1186/1471-2180-11-241/TABLES/3.

[14] Ng EY, Trucksis M, Hooper DC. Quinolone resistance mutations in topoisomerase IV: relationship to the flqA locus and genetic evidence that topoisomerase IV is the primary target and DNA gyrase is the secondary target of fluoroquinolones in Staphylococcus aureus. Antimicrob Agents Chemother 1996;40:1881–8. https://doi.org/10.1128/AAC.40.8.1881.

[15] Takahata M, Yonezawa M, Kurose S, Futakuchi N, Matsubara N, Watanabe Y, et al. Mutations in the gyrA and grlA genes of quinolone-resistant clinical isolates of methicillin-resistant Staphylococcus aureus. J Antimicrob Chemother 1996;38:543–6. https://doi.org/10.1093/JAC/38.3.543.

[16] Avrain L, Garvey M, Mesaros N, Glupczynski Y, Mingeot-Leclercq MP, Piddock LJV, et al. Selection of quinolone resistance in Streptococcus pneumoniae exposed in vitro to subinhibitory drug concentrations. J Antimicrob Chemother 2007;60:965–72. https://doi.org/10.1093/JAC/DKM292.

[17] DelVecchio VG, Petroziello JM, Gress MJ, McCleskey FK, Melcher GP, Crouch HK, et al. Molecular genotyping of methicillin-resistant Staphylococcus aureus via fluorophore-enhanced repetitive-sequence PCR. J Clin Microbiol 1995;33:2141–4. https://doi.org/10.1128/JCM.33.8.2141-2144.1995.

[18] Griggs DJ, Marona H, Piddock LJV. Selection of moxifloxacin-resistant Staphylococcus aureus compared with five other fluoroquinolones. J Antimicrob Chemother 2003;51:1403–7. https://doi.org/10.1093/JAC/DKG241.

[19] Riedel G, Rüdrich U, Fekete-Drimusz N, Manns MP, Vondran FWR, Bock M. An Extended ΔCT-Method Facilitating Normalisation with Multiple Reference Genes Suited for Quantitative RT-PCR Analyses of Human Hepatocyte-Like Cells. PLoS One 2014;9. https://doi.org/10.1371/JOURNAL.PONE.0093031.

[20] Costa SS, Viveiros M, Rosato AE, Melo-Cristino J, Couto I. Impact of efflux in the development of multidrug resistance phenotypes in Staphylococcus aureus Clinical microbiology and vaccines. BMC Microbiol 2015;15:1–16. https://doi.org/10.1186/S12866-015-0572-8/FIGS/7.

[21] Couto I, Costa SS, Viveiros M, Martins M, Amaral L. Efflux-mediated response of Staphylococcus aureus exposed to ethidium bromide. J Antimicrob Chemother 2008;62:504–13. https://doi.org/10.1093/JAC/DKN217.

[22] Tuchscherr L, Bischoff M, Lattar SM, Noto Llana M, Pförtner H, Niemann S, et al. Sigma Factor SigB Is Crucial to Mediate Staphylococcus aureus Adaptation during Chronic Infections. PLoS Pathog 2015;11. https://doi.org/10.1371/JOURNAL.PPAT.1004870.

[23] Miller HK, Carroll RK, Burda WN, Krute CN, Davenport JE, Shaw LN. The extracytoplasmic function sigma factor σS protects against both intracellular and extracytoplasmic stresses in Staphylococcus aureus. J Bacteriol 2012;194:4342–54. https://doi.org/10.1128/JB.00484-12.

[24] Kumar A, Lorand D. Robust ΔΔct estimate. Genomics 2021;113:420–7. https://doi.org/10.1016/J.YGENO.2020.12.009.

[25] Chenia HY, Pillay B, Pillay D. Analysis of the mechanisms of fluoroquinolone resistance in urinary tract pathogens. J Antimicrob Chemother 2006;58:1274–8. https://doi.org/10.1093/JAC/DKL404.

[26] Ruiz J. Mechanisms of resistance to quinolones: target alterations, decreased accumulation and DNA gyrase protection. J Antimicrob Chemother 2003;51:1109–17. https://doi.org/10.1093/JAC/DKG222.

[27] Drago L, Nicola L, Mattina R, de Vecchi E. In vitro selection of resistance in Escherichia coli and Klebsiella spp. at in vivo fluoroquinolone concentrations. BMC Microbiol 2010;10. https://doi.org/10.1186/1471-2180-10-119.

[28] Ince D, Hooper DC. Mechanisms and frequency of resistance to gatifloxacin in comparison to AM-1121 and ciprofloxacin in Staphylococcus aureus. Antimicrob Agents Chemother 2001;45:2755–64. https://doi.org/10.1128/AAC.45.10.2755-2764.2001.

[29] Oyamada Y, Ito H, Fujimoto K, Asada R, Niga T, Okamoto R, et al. Combination of known and unknown mechanisms confers high-level resistance to fluoroquinolones in Enterococcus faecium. J Med Microbiol 2006;55:729–36. https://doi.org/10.1099/JMM.0.46303-0.

[30] Andersson DI, Levin BR. The biological cost of antibiotic resistance. Curr Opin Microbiol 1999;2:489–93. https://doi.org/10.1016/S1369-5274(99)00005-3.

[31] Costa SS, Viveiros M, Amaral L, Couto I. Multidrug Efflux Pumps in Staphylococcus aureus: an Update. Open Microbiol J 2013;7:59–71. https://doi.org/10.2174/1874285801307010059.

[32] Huet AA, Raygada JL, Mendiratta K, Seo SM, Kaatz GW. Multidrug efflux pump overexpression in Staphylococcus aureus after single and multiple in vitro exposures to biocides and dyes. Microbiology (Reading) 2008;154:3144–53. https://doi.org/10.1099/MIC.0.2008/021188-0.

[33] Ding Y, Onodera Y, Lee JC, Hooper DC. NorB, an efflux pump in Staphylococcus aureus strain MW2, contributes to bacterial fitness in abscesses. J Bacteriol 2008;190:7123–9. https://doi.org/10.1128/JB.00655-08.

[34] Thai VC, Lim TK, L. KPU, Lin Q, Nguyen TTH. iTRAQ-based proteome analysis of fluoroquinolone-resistant Staphylococcus aureus. J Glob Antimicrob Resist 2017;8:82–9. https://doi.org/10.1016/J.JGAR.2016.11.003.

[35] Drazic A, Myklebust LM, Ree R, Arnesen T. The world of protein acetylation. Biochimica et Biophysica Acta (BBA) - Proteins and Proteomics 2016;1864:1372–401. https://doi.org/10.1016/J.BBAPAP.2016.06.007.

[36] Lee E ji, Seo JH, Kim KW. Special issue on protein acetylation: from molecular modification to human disease. Experimental & Molecular Medicine 2018 50:7 2018;50:1–2. https://doi.org/10.1038/s12276-018-0103-4.

[37] Christensen DG, Xie X, Basisty N, Byrnes J, McSweeney S, Schilling B, et al. Post-translational Protein Acetylation: An Elegant Mechanism for Bacteria to Dynamically Regulate Metabolic Functions. Front Microbiol 2019;10. https://doi.org/10.3389/FMICB.2019.01604.

[38] Pletnev PI, Shulenina O, Evfratov S, Treshin V, Subach MF, Serebryakova M v., et al. Ribosomal protein S18 acetyltransferase RimI is responsible for the acetylation of elongation factor Tu. J Biol Chem 2022;298. https://doi.org/10.1016/J.JBC.2022.101914.

[39] rimI - [Ribosomal protein S18]-alanine N-acetyltransferase - Escherichia coli (strain K12) | UniProtKB | UniProt n.d. https://www.uniprot.org/uniprotkb/P0A944/entry (accessed November 19, 2022).

[40] Bai J, Zhu X, Zhao K, Yan Y, Xu T, Wang J, et al. The role of ArlRS in regulating oxacillin susceptibility in methicillin-resistant Staphylococcus aureus indicates it is a potential target for antimicrobial resistance breakers. Emerg Microbes Infect 2019;8:503. https://doi.org/10.1080/22221751.2019.1595984.

[41] Stapleton PD, Taylor PW. Methicillin resistance in Staphylococcus aureus: mechanisms and modulation. Sci Prog 2002;85:57. https://doi.org/10.3184/003685002783238870.

[42] Komatsuzawa H, Ohta K, Sugai M, Fujiwara T, Glanzmann P, Berger-Bächi B, et al. Tn551-mediated insertional inactivation of the fmtB gene encoding a cell wall-associated protein abolishes methicillin resistance in Staphylococcus aureus. Journal of Antimicrobial Chemotherapy 2000;45:421–31. https://doi.org/10.1093/JAC/45.4.421.

[43] Maslowska KH, Makiela-Dzbenska K, Fijalkowska IJ. The SOS system: A complex and tightly regulated response to DNA damage. Environ Mol Mutagen 2019;60:368. https://doi.org/10.1002/EM.22267.

[44] Grinholc M, Rodziewicz A, Forys K, Rapacka-Zdonczyk A, Kawiak A, Domachowska A, et al. Fine-tuning recA expression in Staphylococcus aureus for antimicrobial photoinactivation: importance of photo-induced DNA damage in the photoinactivation mechanism. Appl Microbiol Biotechnol 2015;99:9161. https://doi.org/10.1007/S00253-015-6863-Z.

[45] Luong TT, Dunman PM, Murphy E, Projan SJ, Lee CY. Transcription Profiling of the mgrA Regulon in Staphylococcus aureus. J Bacteriol 2006;188:1899. https://doi.org/10.1128/JB.188.5.1899-1910.2006.

[46] Jiang Q, Jin Z, Sun B. MgrA Negatively Regulates Biofilm Formation and Detachment by Repressing the Expression of psm Operons in Staphylococcus aureus. Appl Environ Microbiol 2018;84. https://doi.org/10.1128/AEM.01008-18.

[47] Vogel C, de Sousa Abreu R, Ko D, Le SY, Shapiro BA, Burns SC, et al. Sequence signatures and mRNA concentration can explain two-thirds of protein abundance variation in a human cell line. Mol Syst Biol 2010;6. https://doi.org/10.1038/MSB.2010.59.

[48] Kozak M. Point mutations define a sequence flanking the AUG initiator codon that modulates translation by eukaryotic ribosomes. Cell 1986;44:283–92. https://doi.org/10.1016/0092-8674(86)90762-2.

[49] Lackner DH, Beilharz TH, Marguerat S, Mata J, Watt S, Schubert F, et al. A network of multiple regulatory layers shapes gene expression in fission yeast. Mol Cell 2007;26:145–55. https://doi.org/10.1016/J.MOLCEL.2007.03.002.

[50] Calvo SE, Pagliarini DJ, Mootha VK. Upstream open reading frames cause widespread reduction of protein expression and are polymorphic among humans. Proc Natl Acad Sci U S A 2009;106:7507–12. https://doi.org/10.1073/PNAS.0810916106.

[51] Mayr C, Bartel DP. Widespread shortening of 3’UTRs by alternative cleavage and polyadenylation activates oncogenes in cancer cells. Cell 2009;138:673–84. https://doi.org/10.1016/J.CELL.2009.06.016.

[52] Courel M, Clément Y, Bossevain C, Foretek D, Cruchez OV, Yi Z, et al. GC content shapes mRNA storage and decay in human cells. Elife 2019;8. https://doi.org/10.7554/ELIFE.49708.

[53] Litterman AJ, Kageyama R, le Tonqueze O, Zhao W, Gagnon JD, Goodarzi H, et al. A massively parallel 3’ UTR reporter assay reveals relationships between nucleotide content, sequence conservation, and mRNA destabilization. Genome Res 2019;29:896–906. https://doi.org/10.1101/GR.242552.118.

[54] Mordstein C, Savisaar R, Young RS, Bazile J, Talmane L, Luft J, et al. Codon Usage and Splicing Jointly Influence mRNA Localization. Cell Syst 2020;10:351-362.e8. https://doi.org/10.1016/J.CELS.2020.03.001.

[55] Mauger DM, Joseph Cabral B, Presnyak V, Su S v., Reid DW, Goodman B, et al. mRNA structure regulates protein expression through changes in functional half-life. Proc Natl Acad Sci U S A 2019;116:24075–83. https://doi.org/10.1073/PNAS.1908052116.

